# Individual behavioral signatures shape processing of context-dependent social fear

**DOI:** 10.64898/2026.10.05.754998

**Authors:** Lito Parapera Papantoniou, Alessandro Contestabile, Pierre-Hugues Prouvot Bouvier, Chiara Bezzola, Mattia Lucchini, Corinne Pfister, Orysia Vityk, Balsam Alazmi, Stamatina Tzanoulinou

**Affiliations:** Department of Biomedical Sciences, Faculty of Biology and Medicine, University of Lausanne, Rue du Bugnon 7, 1005, Lausanne, Switzerland; Department of Neuroscience, Karolinska Institutet, Biomedicum, Solnavägen 9, 171 65 Stockholm, Sweden; Department of Fundamental Neurosciences, Faculty of Biology and Medicine, University of Lausanne, Rue du Bugnon 9, 1005 Lausanne, Switzerland; Institute of Complementary and Integrative Medicine, University of Bern, Fabrikstrasse 8, 3012 Bern, Switzerland

**Keywords:** individual differences, contextual fear processing, social defeat stress, conditioned place aversion, VTA, GABA

## Abstract

Context processing is an essential function in the cognitive repertoire of numerous species, including humans and mice, allowing them to learn from their environment and adapt their future behavior. Impairments or alterations in context processing are hypothesized to represent one of the core etiological mechanisms underlying trauma-and stressor-related disorders such as post-traumatic stress disorder (PTSD). Individual behavioral traits are key determinants of post-stress responses; however, how they influence distinct phases of contextual encoding and processing remains incompletely understood. This knowledge gap is further exacerbated by the limited availability of contextual fear protocols based on social stress, even though the latter is among the most frequent and impactful forms of stress exposure. Here, based on social defeat paradigms, we establish a longitudinal social conditioned place aversion (sCPA) protocol that leads to avoidance of the defeat-associated chamber in adult male mice. Prior characterization of mice according to anxiety-related phenotypes revealed that mice displaying high novelty-induced thigmotaxic (*i.e.* high anxiety-like) behavior showed lasting long-term memory of the defeat-associated chamber as denoted by persistent avoidance. On the other hand, lower levels of exploration/locomotor behaviors prior to stress exposure predicted greater avoidance of the defeat-associated chamber in the short-term. Finally, we reveal that inhibiting the activity of VTA^Vgat^ neurons during conditioning sessions abolished sCPA acquisition, while disrupting the predictive ability of locomotor/exploratory profiles derived prior to the sCPA. Our data establishes a robust temporally structured social stress-based conditioning protocol and reveals how prior individual anxiety-and exploration/locomotion-related traits influence distinct stages of context processing.

## Introduction

Contextual fear processing is considered of prime importance in the pathophysiology and symptomatology of multiple psychiatric conditions such as depression, schizophrenia and stress-related disorders.^1^ In particular, posttraumatic stress disorder (PTSD) is categorized as a trauma-and stressor-related disorder according to the DSM-V for which exposure to a traumatic event is a necessary precondition.^2^ Key symptoms of PTSD involve context processing alterations, such as re-experiencing of the stressful event (flashbacks), intrusive memories, hyperarousal and nightmares, with evidence suggesting deficits in fear extinction and impaired retention of extinction.^1,3–6^ Prevalence estimates for stress-related disorders vary widely across sociodemographic groups, which makes precise global figures difficult to establish, yet these mental disorders are consistently recognized as imposing a substantial burden on both individuals and society.^7,8^ Despite important advances that have admittedly increased our understanding of stress-related, anxiety and mood disorders, treatments for these patients remain limited and often ineffective.^9,10^

Not everyone that experiences stressful events will develop psychopathology. This implies that there are inter-individual differences in stress responses that contribute to the extent of maladaptations after a traumatic event.^11–14^ Anxiety-related behaviors, and specifically high anxiety traits, have been linked with vulnerability to stress responses, potentially predisposing the individual to develop psychopathology.^14,15^ Anxious individuals seem to be hyper-responsive to threatening or aversive stimuli, catastrophize more often and focus on worst-case outcomes; features that may be linked to enhanced probability of adopting avoidance strategies.^16–21^ In fact, PTSD was previously categorized as an anxiety disorder, with a subset of clinicians noting that its declassification from anxiety to stressor-related disorder in DSM-V has been detrimental for diagnosis, as anxiety is considered a core hallmark for its development.^22^ Exploration seems to be another crucial factor contributing to contextual fear processing.^23,24^ Using elegant behavioral paradigms, it was shown that mice need to spatially explore the context -in which later they will experience a stressful stimulus -in order to acquire contextual fear memories efficiently.^23^ However, the precise contribution of anxiety traits and exploratory behaviors in specific phases of context encoding and processing is not yet fully mapped, with clear and direct translational implications.

Fear conditioning protocols in rodents, whereby the associations between specific environmental cues and certain experiences are studied, have been an instrumental tool to model features of stress-related disorders.^1,25^ Most rodent contextual fear models involve protocols employing electric shocks as stressors (*i.e.* unconditioned stimuli; US)^25^ and freezing responses (*i.e*. immobility apart from the movements related to breathing) as fear learning and memory proxies.^26^ However, limitations of such models have been highlighted^25^ while widely-used methodologies are being refined. Thus, it is becoming evident that conclusions drawn from electric shock-based fear conditioning protocols are not always applicable to fear learning based on other types of stressors, such as social stress. Regarding fear metrics, it was recently demonstrated that velocity is more relevant than time spent in proximity to an aggressor in denoting social avoidance, particularly in females^27^. Moreover, studies have clearly revealed non-overlapping brain circuitry between learned social stress and electric shocks or innate stressors such as predator odors,^28–30^ suggesting that the type of stress itself can be crucial in determining the underlying neurobiology. Despite social stress being an ethologically relevant stressor for social species such as rodents and humans,^13^ paradigms of contextual fear learning based on social stress remain but a few.^29,31–33^ Therefore, expanding contextual fear paradigms to include other types of stress and fear proxies, will aid to map contributions of inter-individual differences to stress response vulnerability, dissect the underlying neurobiology, and thus, significantly advance the field.

While a significant volume of research has documented the role of brain areas such as the hippocampus, the prefrontal cortex and the amygdala in contextual fear processing,^1,25^ recent studies have revealed key roles of other areas, such as hypothalamic subregions^28,32^ and the ventral tegmental area (VTA)^31,34^ in rodent social stress paradigms. Previously, it was shown that mice exposed to an acute stressor showed an increased plasticity in VTA dopamine (DA) neurons, which could be blocked by a glucocorticoid receptor (GR) antagonist.^35^ These neurons also showed increased synaptic strength during the acquisition of an associative learning task for a reward-predicting cue (but not once behavioral performance is stable).^36^ On the other hand, VTA GABA neurons are critical for conditioned place aversion^37^ and are shown to be activated upon social defeat stress, with somatostatin (SST) neurons highlighted as a key subpopulation.^38^ Despite advances denoting the participation of VTA cell populations in reward and aversion,^31,39^ the mechanistic role of specific VTA neuronal populations to distinct context encoding and conditioning phases in paradigms of contextual social fear learning remains incompletely understood.

Based on protocols of social defeat stress,^13,40,41^ we first establish and characterize a novel longitudinal and temporally-structured conditioned place aversion task in male mice allowing us to probe distinct context encoding, conditioning and retention phases with precise temporal and spatial resolution. Next, we employ machine learning tools for automatically scoring individual behaviors to identify potential new proxies of fear responses and ask whether prior inter-individual differences in anxiety-like and exploratory behaviors can be associated with contextual social fear processing. Finally, using immunohistochemistry and cre-dependent chemogenetic manipulations, we enquire about the underlying brain areas in our task. Although this work does not aim to model PTSD or specific stress-related disorders, we unravel key features of social fear context processing, which will in turn help to advance translational approaches.

## Materials and Methods

### Animals

All experimental procedures described here were performed in accordance with the Swiss laws and were previously approved by the Swiss Cantonal Veterinary Office Committee for Animal Experimentation. Experimental C57BL/6J (BL6) male mice were purchased from Charles River Laboratories and were group-housed (2 per cage) in the institutional animal facility under standard 12h/12h light/dark cycle with food and water *ad libitum*. Experimental B6J.129S6(FVB)-Slc32a1tm2(cre)Lowl/MwarJ (Vgat-cre) male mice were also group-housed (2-4 per cage) in the institutional animal facility under standard 12h/12h light/dark cycle with food and water *ad libitum.* All experimental mice were tested in the behavioral experiments described below during adulthood. Behavioral experiments were performed during the light phase (between 9 am and 5 pm) in a room with dim light conditions and temperature between 22 and 24°C. CD-1 male retired breeders (4-9 months old) were purchased from Charles River Laboratories and were single-housed in the institutional animal facility under standard 12h/12h light/dark cycle with food and water *ad libitum.* CD-1 mice were housed in a separate room from the experimental mice to avoid sensory cues transfer. BL6 male juvenile mice (around postnatal day 27 -P27) that were used as stimuli mice in the Social Preference test were purchased from Charles River Laboratories and were group-housed (2-4 per cage) in the institutional animal facility under standard 12h/12h light/dark cycle with food and water *ad libitum*. At the end of each experiment, all mice returned to their home cages.

### Screening of aggressor CD-1 mice

The standard procedure as described by Golden et al. (2011) was followed.^40^ The procedure expands over three days. Each day, one young BL6 male screener mouse (8-9 weeks old) was introduced in the home cage of a CD-1 mouse for 180 seconds, and the latency to receive an attack by the CD-1 mouse was noted. Each CD-1 mouse should encounter a different screener mouse every day. Only CD-1 mice with attack latencies of no more than 60 seconds per day, for at least two consecutive days, were considered to have passed the test and could be used as aggressors.

### Handling

Prior to any behavioral experiment, experimental BL6 and Vgat-cre mice were subjected to three handling sessions of 90 seconds each, spanning over 2-3 days. Vgat-cre mice were habituated to systemic injections with one intraperitoneal injection (i.p.) of saline daily during the days of handling.

### Behavioral testing

All behavioral experiments, including social Conditioned Place Aversion (sCPA), Open Field (OF), reactivity to a Novel Object (NO), Elevated Plus Maze (EPM), Novel Object Recognition (NOR), 3-Chamber Social Preference (SP), and CD-1 Social Interaction (CD-1 SI) tests, were video recorded and tracked using EthoVision XT software versions 15 and 16 (Noldus Information Technology, Wageningen, the Netherlands). The order of behavioral testing was kept consistent across animals within each concerned experimental cohort and was conducted in the following sequence: OF, NO, EPM, sCPA, OF, NO, NOR, SP, CD-1 SI. Between animals, the apparatuses were cleaned with 5% ethanol solution.

### Social Conditioned Place Aversion (sCPA)

The social Conditioned Place Aversion (sCPA) protocol was performed in a commercially available apparatus (Bioseb, Model BIOSEB, in vivo Research Instruments, spatial place preference box for mice LE895).^42^ The apparatus consists of two square-shaped chambers (20 x 20 cm) that communicate through a small rectangular transparent corridor. The two chambers contain different visual patterns, with either black dots or grey stripes on white background on the walls. Each chamber communicates with the corridor through a removable door. Experimental mice were randomly assigned to one of the following categories: control (object-based conditioning) or aversive social conditioning (sCPA). The protocol for the sCPA group is described below (**Fig. 1A**).

**Figure 1 -.**
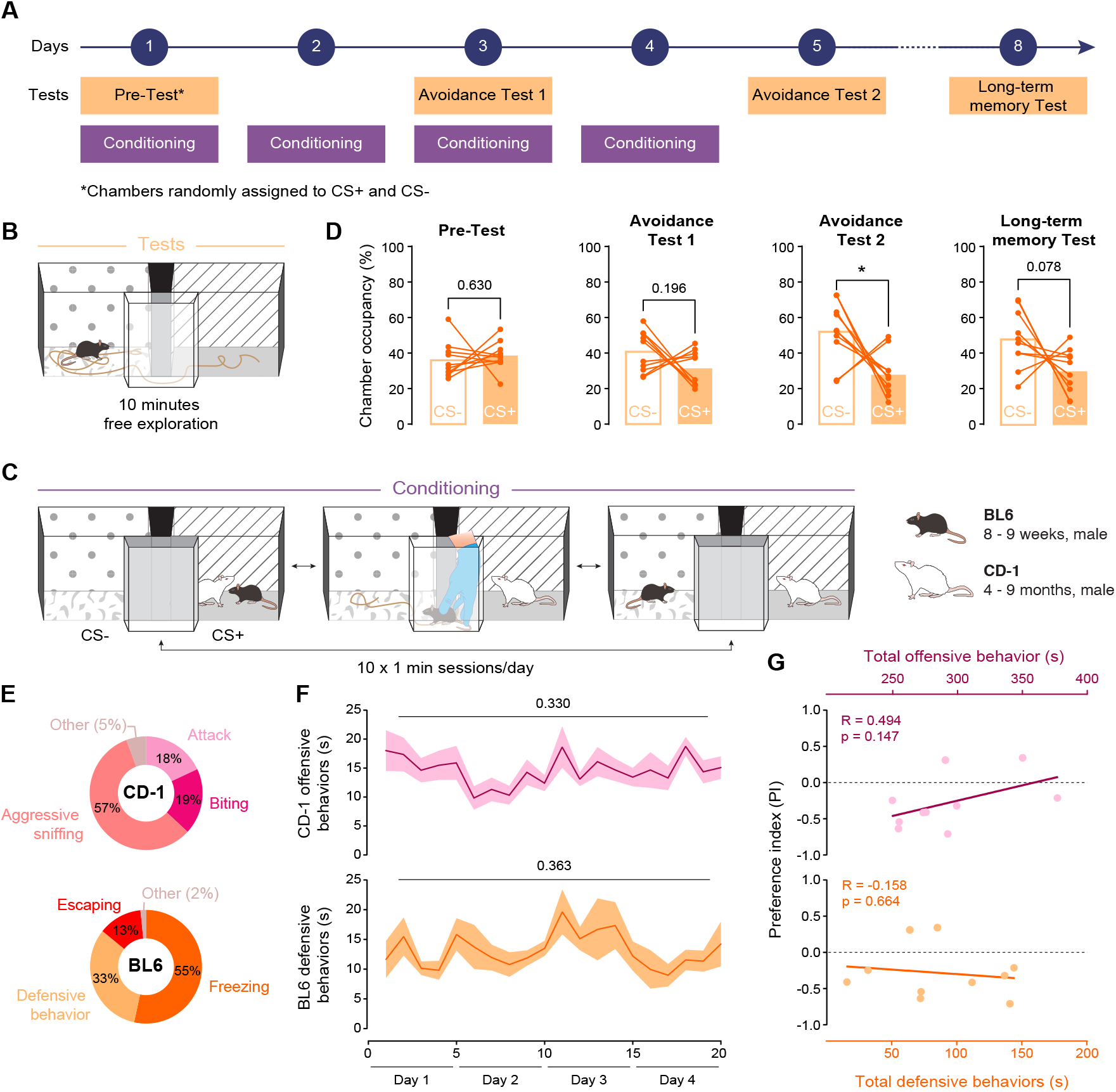
A social Conditioned Place Aversion paradigm reveals avoidance of an aggressor-paired context. **A**, Experimental design. Mice underwent a Pre-Test session followed by four days of conditioning. Avoidance was assessed after two and four conditioning sessions (Avoidance Test 1 & 2, respectively). Three days later, individuals’ preference/avoidance of the chambers was assessed via a Long-term memory Test. During the Pre-Test, chamber identities were randomly assigned as conditioned stimulus positive (CS+) and conditioned stimulus negative (CS−). **B**, Testing procedure. Mice freely explored the apparatus for 10 minutes (min), and occupancy of the two chambers was quantified. **C**, Conditioning procedure. During conditioning, BL6 mice were confined either with an aggressive CD-1 mouse in the CS+ chamber or alone in the CS− chamber in alternating sessions (10 ✕ 1-min sessions per day). **D**, Occupancy of the CS− and CS+ chambers during the Pre-Test, Avoidance Test 1, Avoidance-Test 2, and Long-term memory Test. Each line represents an individual animal. Pre-Test (paired t-test): t(9) = −0.499, p = 0.630; Avoidance Test 1 (paired t-test): t(9) = 1.398, p = 0.196; Avoidance Test 2 (paired t-test): t(9) = 2.631, p = 0.027; Long-term memory Test (paired t-test): t(9) = 1.988, p = 0.078. **E**, Distribution of offensive and defensive behaviors in percentage of total scored behaviors displayed by CD-1 aggressors and BL6 mice during conditioning. **F**, Temporal dynamics of CD-1 offensive behaviors and BL6 defensive responses across conditioning sessions. Shaded areas represent mean ± SEM. CD-1 (RM one-way ANOVA with Greenhouse-Geisser correction): F(5.190, 46.707) = 1.187, p = 0.330; BL6 (RM one-way ANOVA with Greenhouse-Geisser correction): F(5.789, 52.105) = 1.121, p = 0.363. **G**, Relationship between total offensive behavior experienced (top) or total defensive behavior exerted (bottom) during conditioning and avoidance expressed during Avoidance Test 2. Each point represents one animal. Relationship between total offensive behavior and avoidance, Statistical Test: Pearson’s correlation, R = 0.494, p = 0.147. Relationship between total defensive behavior and avoidance: Statistical test: Pearson’s correlation, R = −0.158, p = 0.664.

#### Pre-Test (Day 1)

During the first day of the protocol, each experimental mouse was placed in the apparatus with the doors open and was left for 10 minutes to freely explore.

#### Conditioning Days (Days 1-4)

In total, four conditioning days take place, during which each experimental mouse encounters a CD-1 mouse in the conditioned chamber (CS+ chamber), whereas the other chamber of the apparatus is empty and thus, referred to as the unconditioned chamber (CS-chamber). In total, two CD-1 mice were used in alteration for each experimental mouse throughout the conditioning days. The first conditioning session takes place right after the Pre-Test is completed. To eliminate potential biases, mice were randomly assigned to a CS+ chamber, either the one with dots, or the one with the stripes wall pattern. Each conditioning day started with placing each mouse in one of the two chambers (CS+ or CS-) and consisted of 9 alterations between them. Experimental mice were restricted to each chamber for 60 seconds with the doors closed every time (10 conditioning trials in total; 5 in the CS+ and 5 in the CS-). At the end of each conditioning trial, the doors were lifted and the experimental mice were gently guided to the other chamber. The order of chamber alterations was counterbalanced between days for each experimental mouse. For example, if a given mouse starts in the CS+ chamber on the first conditioning day, on the second day, it starts in the CS-chamber and so on. To avoid injuries, in cases of excessive aggression, the trial was interrupted and the BL6 mouse was gently guided in the CS-chamber until the end of the CS+ session. For the remaining within-day conditioning sessions of that mouse, a wire-mesh enclosure containing the aggressive CD-1 mouse was placed in the CS+ chamber to prevent wounding of the BL6, and the conditioning trials continued normally.

#### Avoidance Test 1 (Day 3)

The Avoidance-Test 1 is performed 24 hours after the second conditioning day and precedes the third conditioning session. Each experimental mouse was placed in the apparatus with the doors open in the absence of the CD-1 mouse and was left to freely explore it for 10 minutes.

#### Avoidance Test 2 (Day 5)

24 hours following the fourth conditioning day, each experimental mouse was placed in the apparatus with the doors open in the absence of the CD-1 mouse and was left to freely explore it for 10 minutes.

#### Long-term memory Test (Day 8)

72 hours after the Avoidance Test 2, each experimental mouse was again placed in the apparatus with the doors open in the absence of the CD-1 mouse and was left for a 10-minute free exploration.

The control group underwent the same procedure but instead of a CD-1 mouse, each experimental mouse encountered a neutral object (small metallic locker) in the CS+ chamber during the conditioning sessions (object-based conditioning).

Regarding chemogenetic inhibition of Vesicular GABA Transporter (Vgat) neurons in the VTA during conditioning, Vgat-cre mice were i.p.-injected with 0.3 mg/kg clozapine-N-oxide (CNO) 30-40 minutes before the first conditioning trial of each conditioning day. No injections were performed before test sessions. It is noted that one hM4Di-injected mouse was not detected during the first 2.5 minutes of the Avoidance-Test 1 due to occlusion while being at the CD-1 chamber, therefore chamber occupancy for this bin was manually set to 100% in the observed chamber following video inspection.

### Open Field (OF) – reactivity to a Novel Object (NO) tests

Locomotion and anxiety-like behavior were evaluated as previously described in open field square-shaped arenas (Semadeni, sides 45 cm ✕ height 40 cm) with the Open Field (OF) test. ^43,44^ Each experimental BL6 mouse was introduced in the arena and was left to freely explore it for 10 minutes. The incentive to explore a novel object in an anxiogenic setting was assessed in a subsequent phase with the reactivity to a Novel Object (NO) test during which, a novel object (small metallic locker) was introduced in the center of the arena and the mouse was left to explore freely for 5 minutes. Distance traveled, exploration time, and frequency of exploration episodes for the center (zone including the object) and walls of the arena were assessed.

### Elevated Plus Maze (EPM) test

Anxiety-like behavior was also evaluated with the Elevated Plus Maze (EPM) test inside a cross-shaped apparatus with two open and two closed arms (arms’ dimensions: 30 cm long ✕ 5.5 cm wide) elevated 56 cm from the floor.^43,45^ Each experimental BL6 mouse was placed in the center of the apparatus and was left to freely explore for 5 minutes. Distance traveled, exploration time, and frequency of exploration episodes for the center, open, and closed arms were measured.

### Novel Object Recognition (NOR) test

During the habituation day of the test, each experimental BL6 mouse was placed twice in the OF arenas for 5 minutes each time to freely explore, with a 30 minute-long interval. The following day, two identical objects (height 16 cm) were placed adjacent to one another close to the wall. The experimental mouse was introduced to the arena and was left to explore for 10 minutes. The next day, each mouse was placed again in the objects-containing arena, while this time, one of the previously explored (familiar) objects was replaced with a novel object of similar height and each mouse was left to freely explore for 10 minutes. Exploration time and frequency of exploration episodes were manually scored with EthoVision XT by an experimenter blind to the experimental conditions. The novel object exploration ratio was calculated as follows:

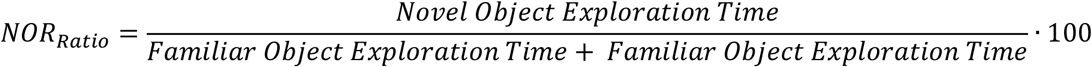

### 3-Chamber Social Preference (SP) test

Sociability was assessed as previously described.^46,47^ The 3-chamber social preference (SP) test was performed in a rectangular Plexiglass arena (60 x 40 x 30 cm). The apparatus is divided into three equally-sized rectangular chambers (40 x 20 x 30 cm) that communicate through doors that can be lifted to provide access to all chambers. Vgat-cre mice were administered i.p. with CNO 30 minutes before the SP test. During the habituation phase of the test, each experimental mouse was placed in the central chamber with the doors open and was left to freely explore the arena for 10 minutes. At the end of the habituation phase, the mouse was gently guided into the central chamber and the doors closed to restrict it there for a few seconds while the experimenter placed an enclosure (diameter 9 cm ✕ height 18 cm) in the center of each of the other two chambers. One enclosure was empty and the other one contained a non-familiar juvenile conspecific. The doors were then lifted and the experimental mouse was left to freely explore for 10 minutes. Juvenile mice had already been habituated to the enclosures with three 10 minute-long sessions a few days before the SP test. Distance traveled, mean velocity, exploration time, frequency of exploration episodes around the juvenile-containing and empty enclosures were measured. The juvenile exploration ratio was calculated as follows:

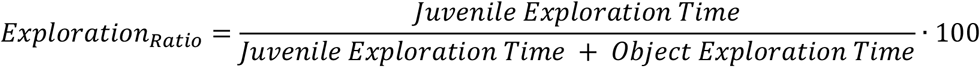

One animal from the mCherry-expressing group was excluded from the analysis as it showed no exploration for the juvenile mouse.

### CD-1 Social Interaction (SI) test

To assess the long-term memory for the CD-1 mice, experimental BL6 mice underwent the CD-1 Social Interaction (CD-1 SI) test as previously described.^40,41^ During the first phase of the experiment, an empty wire-mesh enclosure was placed in close proximity to one wall of the OF arena and the experimental mouse was allowed to freely explore the apparatus for 2.5 minutes. During the second phase, the empty enclosure was replaced by an enclosure containing a novel CD-1 mouse and the experimental mouse was allowed to freely explore for an additional 2.5 minutes. Exploration time in the interaction zone (IZ) and corners, in the presence (+) or absence (-) of the CD-1 mouse, were measured. The CD-1 SI Ratio was calculated as follows: Time in the IZ with a CD-1 (+) (s) / Time in the IZ without a CD-1 (-) (s). Two animals, one from the object-based conditioning group and one from the sCPA group, were excluded from the SI ratio analysis as outliers, with values outside the expected range.

### Manual scoring of social defeat

Offensive behaviors of the CD-1 mice, as well as behavior of the experimental BL6 mice during social defeat were scored manually using EthoVision XT (**Fig. 1**, **Fig. S1**). Regarding CD-1 offensive behaviors, we measured the duration of attacks, biting, aggressive sniffing, offensive upright, lateral threat and keeping down episodes across days. We grouped all these behaviors as an “offensive behavior” composite (**Fig. 1E-F**). For clarity, we grouped offensive upright, lateral threat and keeping down, that exhibited minimal duration, under the category “Other” in the pie chart presented in **Fig. 1E** (top). Pertaining to BL6 behavior, we measured the duration of freezing, defensive behavior, escaping, submissive behavior, rearing, and self-grooming across days. Similarly, we grouped submissive behavior, rearing, and self-grooming, that exhibited minimal duration, under the category “Other” in the pie chart presented in **Fig. 1E** (bottom). Pie charts in **Fig. 1E** present percentages corresponding to subdivisions of the total scored behavior for each group. Regarding the temporal dynamics presented in **Fig. S1**, the mean percentage of each behavioral subdivision and corresponding SEM were plotted across conditioning trials.

### Identification of behavioral subgroups based on thigmotaxis

To determine whether animals could be separated into distinct behavioral subgroups based on thigmotaxis, the distribution of time spent in the wall zone (% of total session time) during reactivity to a Novel Object (NO) was modeled using a two-component Gaussian mixture model. Model parameters were estimated by maximum likelihood, and the two mixture components were interpreted as low-and high-anxiety (*i.e.* low-and high-thigmotaxis respectively) populations according to their estimated means. A classification threshold was defined as the point at which the weighted probability density functions of the two fitted Gaussian components intersected, corresponding to an equal likelihood of assignment to either population. Animals with values below this threshold were classified as belonging to the low-anxiety group, whereas animals with values above the threshold were classified as belonging to the high-anxiety group. For visualization, the empirical distribution of thigmotaxis values was represented using kernel density estimation, and individual animals were overlaid as jittered points (**Fig. 3B**).

### Behavioral detection and segmentation

A proprietary deep neural network (DNN) model was trained within the Tomb-AI platform managed by Metofico LTD. (Metofico LTD. Manchester, United Kingdom) to detect the following behaviors: freezing, rearing, grooming, sniffing and a negative category. A database of behavioral events was curated across 94 behavioral videos from various contexts (**Supplementary Table 1**). First, the different behavioral categories were defined as video sequences scored using the Behavioral Observation Research Interactive Software (BORIS University of Turin, Italy). Then precise criteria with an emphasis on beginning and end frames were defined as follows:

#### Freezing

Complete behavioral arrest and immobility to the exclusion of breathing movements. Snout movements such as sniffing and whisking are excluding criteria. Short interruption between freezes was excluded. Freezing occurring during other categories such as rearing is scored as freezing. *Start and end frames:* first and last frames when the animal was completely immobile.

#### Rearing

Animal standing up on its hind legs, lifting the front legs off the floor to explore vertically. *Start frame:* first frame where the front legs have visibly left the floor. If not visible, the first frame when the surface of the visible silhouette of the animal has reduced size and the reflection of light of the coat on the whole back of the animal has shifted to represent the upright position. *End frame:* second frame after the front paws have left the surface on which they were resting. If paws are not visible during the episode, then the first frame when the reflection of the coat on the back of the animal has shifted to return to horizontal position is considered.

#### Grooming

animal rubbing its paws against its body, licking its body or scratching it. Includes face grooms where the paws pass over the face in a postero-anterior manner often after licking them, body grooms where the animal is licking and rubbing its paws against a body part (*i.e*. its rump) periodically and scratches in which the animal claws at its body with periodic movements typically of one hind paw against the shoulders or head. *Start frame:* first frame of the paws passing over the ears or eyes, of the licking or scratching of a body part. If the paws are not visible and periodic movements are unclear or could be mixed up with other behaviors this event is not scored as grooming. Rump grooms and scratches start when one sees the animal bending over 90 degrees. *End frame:* last frame before return to a neutral position or another behavioral category. For scratching, last frame before the body crosses 90 degrees to return to a straight position.

#### Sniffing

Animal performing periodic snout movements and whisking movements to investigate either an object, a wall, the floor or the air. Alternatively, the same snout and whisking movements performed with the head looking up in an almost-rearing position but with all paws on the floor. For most of these events the animal needs to stop its locomotion and have reached null velocity. That is, except in the case of en-passant sniffings where the animal performs visible snout and whisking movements along a surface or an object while being in contact with it, then the animal can exceptionally slow down but not fully stop. *Start frame*: first frame of the head stopped to sniff or of the snout angling in contact with a surface, in the case of sniffing the floor, when the head reaches its end position after coming down and/or retracting. *End frame:* last frame before breaking contact with the object, the wall or the floor. For sniffing with no contact, last frame before retracting snout and head.

#### Negative

any behavior not included in the previous categories, including transition frames in between other categories where the criteria specified above were not reached. Most of the negative behavior consists of locomotion, occlusions, partial or total and periods of unidentified inactivity at rest outside of freezing. During scoring particular emphasis was put on event and positional diversity.

While classical data augmentation was used during training, we also employed a strategy to provide the model with both full-frame spatial context and animal-centered information. All videos were processed using cropping around the arena to maximize the proportion of pixels containing animals while retaining the spatial context of the arena. To further encourage the model to learn animal-centric features, we selected a subset of videos with minimal occlusions and tracked the animals throughout each video. A custom MATLAB script (MATLAB R2020b) was then used to crop each video around the detected center point of the tracked animal, retaining a region corresponding to 20% of the original frame and centered on the animal. This produced animal-centered videos in which the tracked animal remained at the center of the frame. These animal-centered crops were added to the training dataset alongside their corresponding full-frame videos, thereby providing both spatial-context and animal-focused representations of the same behavioral examples. This augmentation strategy was used to train the final model.

Behavioral events were cut into clips of 16 frames, and a first round of training was performed. Clips on the extreme edge of unreliable classification were manually reviewed and removed from the sets if an experienced scorer could not reliably re-classify them, resulting in the support clip numbers in **Supplementary Table 2**. Training clips were two times the number of support clips. After 14 training epochs the model reached a plateau attaining the scores as reported in **Supplementary Table 2**. Frame classification was then performed with Metofico’s proprietary architecture resulting in csv files indicating beginning and ending frames of each event associated with a confidence value. We manually reviewed an out-of-set representative sample of videos from the present publication and empirically defined filtering parameters seen in **Supplementary Table 3**. The filtering parameters were chosen to ensure the highest possible precision at the cost of recall. Precision is defined as *Precision* = (*True Positive* ÷ (*True Positive* + *False positive*)) Recall as *Recall* = (*True positive* ÷ (*True positive* + *False Negative*)) and F1 score as *F1* = *2* × ((*Precision* × *Recall*) ÷ (*Precision* + *Recall*)). Proportion values for each behavior were then calculated using MATLAB. Total time spent performing a behavior was defined as the percentage of time spent performing this behavior either over the whole trial (temporal characterization) or over the time spent in a zone (spatial characterization). The number of episodes was the number of events as defined by each start and end timestamp.

Precision is understood as the accuracy of the model to properly classify a clip, recall the accuracy to detect a behavior and F1-score is the harmonic mean of precision and recall, a common accuracy metric in machine learning. Classification of support clips was used for this accuracy validation. A third of the clips were used for support, bringing the total clip number to 3 times the amount presented here. Please see **Supplementary Video 1** for examples of detected behaviors as the categories presented above.

### Velocity analysis

Animal tracking coordinates and velocities were extracted with no smoothing profile using EthoVision XT. Velocities were then corrected in MATLAB using linear interpolation for missing segments, artifact removal (velocities > 5 ✕ standard deviation) and moving averaging (5 frames window) to remove artifacts due to temporary loss of tracking caused by occlusions.

Pauses were then defined as periods where the velocity of the animal was under 2cm/s and lasting more than 4 seconds; moving segments were defined as the rest of the velocity vectors. Pause duration percentage was calculated as the total duration of Pauses divided by the total duration of the trial. In the case of chamber-based analysis, pauses were quantified as the total time paused in that chamber divided by the total time spent in the corresponding zone. Mean moving velocity was calculated as the mean of the moving velocity excluding the previously defined pauses either over the whole duration of the trial or over the time spent in each zone.

To measure transition velocities we defined transitions as the direct passage from one chamber to the next using the binary “in zone” vectors provided by EthoVision XT. We obtained four kinds of transitions (entries and exits): from CS+ to corridor, from corridor to CS+, from CS-to corridor and from corridor to CS-. We then extracted segments of the pre-processed velocity vector over 24 frames (0.96 s) centered on the transition. We defined the mean transition velocity as the mean of the maximum velocities attained over that segment. Delta velocity was then calculated as such: Delta Velocity = Mean Entry Velocity-Mean Exit velocity.

### Machine learning analysis of behavioral predictors

#### Dataset and preprocessing

All analyses were performed in R (version 4.5.2) using custom scripts. Behavioral features of predictors were derived from Pre-Test sessions and included EthoVision-based measures of kinematics, as well as previously defined behavioral variables using Metofico tomb-AI, capturing both behavioral states and spatial dynamics. Behavioral features included measures of occurrence and frequency of freezing, grooming, rearing, and sniffing, quantified both within the corridor and across the entire apparatus without chamber subdivision. Global locomotor activity was assessed using mean and median velocity, as well as the proportion of time spent paused. Corridor-related variables included total time spent in the corridor, frequency of visits, and visit duration statistics (mean and standard deviation). Movement within the corridor was further quantified as time spent moving and not moving. To capture temporal dynamics, corridor-related measures were also computed across successive time bins (0–150 s, 150–300 s, 300–450 s, and 450–600 s). Variables related to chamber frequency or time spent in chambers were excluded a priori to avoid redundancy and potential circularity with the outcome variable. Predictors with fewer than five non-missing observations or zero variance were removed. The target variable was the Avoidance Test 2 preference index, computed as:

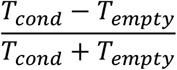

where *cond* refers to the CS+ chamber, and *empty* to the CS-chamber.

#### Model definition and validation

A gradient-boosted decision tree model was trained using the XGBoost library (xgboost_3.2.0.1) to predict Avoidance Test 2 preference from Pre-Test behavioral variables. The model was trained using a learning rate of 0.03, a maximum tree depth of 4, a minimum child weight of 2, subsampling of 0.7, column subsampling of 0.5, and L2 regularization (λ = 1). To limit model complexity and improve generalization, feature selection was performed prior to cross-validation. An initial model was trained on the full set of predictors, and features were ranked based on gain-based importance. The top 20 features were retained for all subsequent analyses. Model performance was assessed using repeated k-fold cross-validation. The dataset was randomly partitioned into five folds, and the model was trained and evaluated across 300 independent repetitions using different random splits. For each repetition, predictions were generated on held-out folds (out-of-fold predictions), ensuring that each animal was evaluated on data not used during training. Performance was quantified using the Pearson correlation coefficient between observed and predicted values as the primary metric. Root mean squared error (RMSE) was also computed to quantify prediction error in the original scale of the target variable, providing a complementary measure of model accuracy. The statistical significance of model performance was assessed using permutation testing. The target variable was randomly permuted 5,000 times, and the full cross-validation procedure was repeated for each permutation. For each iteration, the Pearson correlation between observed and predicted values was computed. An empirical p-value was calculated as:

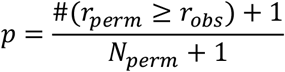

where *r_obs_* is the observed correlation and *r_perm_* represents correlations obtained under permutation. To assess generalizability, the model was evaluated on an independent dataset consisting of animals from a separate experimental cohort, including control virus-injected and Designer Receptors Exclusively Activated by Designer Drugs (DREADD)-injected mice. These animals were excluded from training and used solely for external evaluation.

#### Model interpretation

Feature importance was assessed using gain-based importance scores derived from the trained XGBoost model. To characterize the relationship between individual predictors and model output, SHAP (SHapley Additive exPlanations) values were computed for the top-ranked features, allowing visualization of both the magnitude and direction of the feature.

### Viral injections

For all surgical procedures described hereinafter animals were anesthetized using a mix of oxygen (1 L/min) and isoflurane (3%, Matrx, VIP 3000), they received a subcutaneous injection of carprofen (5 mg/kg) and were placed in a stereotaxic frame (David Kopf Instruments, Model 940). Once the head of the animals was fixed with the help of ear bars, the isoflurane concentration was then reduced and maintained at 1.5%. The skin was then shaved, anesthetized with 50 μL of Lidocaine 0.5% and disinfected with Betadine. After skin incision, craniotomies (1 mm diameter) were performed above the VTA. Injections of Cre-dependent inhibitory DREADD viral vector (ssAAV-5/2-hSyn1-dlox-hM4D(Gi)_mCherry(rev)-dlox-WPRE-hGHp(A), Viral Vector Facility, University of Zurich, v84-5) or the complementary Cre-dependent control vector (ssAAV-5/2-hSyn1-dlox-mCherry(rev)-dlox-WPRE-hGHp(A), Viral Vector Facility, University of Zurich, v116-5) were then performed in Vgat-cre mice (age: 8-10 weeks). Micropipette (Hirschmann, ringcaps® 1-5µl) were inserted in bilateral craniotomies (AP: −3.3mm; ML: ±0.5mm; DV: −4.25mm) and a volume of 200µl was released in the brain tissue over 5 minutes before micropipette retraction.

### c-Fos immunohistochemistry and cell quantification

Sixty minutes after Avoidance Test 1 slices containing key brain regions were collected for c-Fos assessment from sCPA mice. Tissue was also collected from control animals exposed to the object-based conditioning procedure. Coronal brain slices of the whole brain (50 μm thickness) were collected with a vibratome (Leica, VT1200). Brain tissue was rinsed 3 times with PBS 1✕ before incubation with a blocking solution of 10% bovine albumin serum, 0.3% Triton X-100 prepared in PBS 1✕ for 1.5 hours at room temperature. Brain slices containing hippocampal and amygdalar regions were then incubated for 48 hours at 4°C on an orbital shaker with a 3% bovine albumin serum, 0.3% Triton X-100 solution prepared in PBS with a rabbit anti-c-Fos primary antibody (Synaptic Systems, 226 008) diluted at 1:4000. Brain slices containing the VTA were incubated with the same conditions and the same solution but with the addition of a mouse anti-TH primary antibody (MilliporeSigma, MAB318). After incubation with primary antibodies brain slices were rinsed 3 times with PBS 1✕ and then incubated with a 0.25% Tween-20 solution prepared in PBS 1x containing a goat anti-rabbit (Alexa Fluor® 488) secondary antibody (Abcam, ab150077) at a 1:500 dilution for 1.5 hours at room temperature. Slices containing the VTA were also incubated with a goat anti-mouse (Alexa Fluor® 647) secondary antibody (Abcam, ab150115). Brain slices were then rinsed 3 more times with PBS 1✕ before being mounted on glass slides and covered with an antifade mounting medium containing DAPI (Sigma-Aldrich, F6057) and coverslips.

Whole-slides fluorescence images were acquired using an automated slide scanner (Carl Zeiss Microscopy, Axioscan Z1). Image analysis was performed using QuPath (v0.6.0). Prior to cell quantification, section images were registered to a reference atlas (Adult Mouse Brain -Allen Brain Atlas V3p1) using Aligning Big Brains & Atlases (ABBA)^48^ to delineate anatomical brain regions of interest (BLA, LA, VTA, dorsal hippocampus (HPC): CA1, CA2, CA3, DG). Subsequently, quantification of tyrosine hydroxilase-positive (TH⁺) cells was performed using a custom Groovy script in QuPath. In contrast, c-Fos-postive (c-Fos⁺) cells and TH⁺/c-Fos⁺ co-expressing (double-labeled) cells were quantified manually using the built-in annotation and counting tools in QuPath.

### Euthanasia procedures

Four to five weeks after the last defeat session, BL6 experimental mice underwent the CD-1 SI test as described above and were euthanised for tissue and blood collection. Cohorts that were then assigned to immunohistochemistry procedures (BL6 mice for c-Fos assessment and Vgat-cre mice for injection site validation) were transcardially perfused. Specifically, BL6 mice were individually housed for 60 minutes after the CD-1 SI test to avoid unspecific neuronal activation due to external factors, cues, or home cage interactions. Deep anesthesia before perfusion was achieved using a Sodium Pentobarbital i.p injection (250mg/Kg). After ensuring deep anesthesia, the chest cavity was opened to expose the heart and a small needle was inserted into the left ventricle, while the right atrium was cut for drainage. Ice-cold 1✕ PBS was pumped through to flush out the blood, followed by 4% paraformaldehyde (PFA) to fix the tissue.

After a minimum of one week following the last defeat session of the sCPA, Vgat-cre mCherry and mCherry-hM4Di mice were i.p. injected with CNO (0.3mg/kg) and 30 to 40 minutes afterwards exposed to an acute 5 minute-long social defeat session in the home cage of a novel CD-1 mouse. Subsequently, they were individually housed for 30 minutes before being terminally anesthetized with an i.p. administration of pentobarbital as described above. Blood was collected from the heart for corticosterone assessment before the mice were transcardially perfused.

### Corticosterone measurement

Corticosterone measurement was adapted from a procedure previously described.^49^ Blood samples from Vgat-cre mice were collected into heparin-coated capillary tubes (Sarsted, Switzerland, ref.:16.443) and were kept on ice until centrifuged (4,000 rpm at 4°C for 10 minutes). Plasma was collected into new 1.7 ml tubes and stored at -80°C until further analysis. Circulating corticosterone levels were quantified in the plasma samples (1/40 dilution) using an enzymatic immunoassay kit, according to the manufacturer’s guidelines (Enzo Life Sciences, Switzerland, ref.: ADI-900-097). A standard curve method was used to calculate corticosterone levels. It is noted that two mCherry-hM4Di-injected mice were excluded from the analysis as one of them was classified as an outlier and the other one did not yield sufficient blood volume.

### Statistical Analysis

Statistical analysis was conducted with GraphPad Prism (version 11.0.2, San Diego, CA, USA), SPSS (version 29.0.2.0., IBM Corp, 2012), and R (version 4.5.2). The normality of sample distributions was assessed with the Shapiro–Wilk criterion and when violated, non-parametric tests were used. Specifically, when normality was violated, the data were analyzed with Mann–Whitney or Wilcoxon tests, while for multiple comparisons, Kruskal–Wallis or Friedman test was followed by Dunn’s test. When normally distributed, the data were analyzed with independent t-tests, one sample t-tests, one-way ANOVA, and repeated measures (RM) ANOVA as appropriate and stated in the figure legends. When needed, multiple comparisons were adjusted using the Benjamini–Hochberg False Discovery Rate procedure. For the analysis of variance with two factors (two-way ANOVA and RM two-way ANOVA), sphericity was controlled with the Mauchly’s test of sphericity, and the Greenhouse-Geisser correction was considered. Outliers were removed as mentioned in the corresponding methods sections according to the specified criteria. Data are represented as the mean ± SEM and the significance was set at 95% of confidence.

## Results

### Social defeat stress-based conditioning leads to conditioned place aversion

We first sought to develop and characterize a protocol that would allow us to study contextual fear learning, based on social stress. Building upon social defeat protocols, we designed a social Conditioned Place Aversion (sCPA) paradigm that includes distinct test and conditioning sessions across 8 days (**Fig. 1A**). C57BL/6J (BL6) mice were exposed to an arena composed of two separate chambers connected by a corridor to establish their baseline performance (Pre-Test, **Fig. 1B**). This phase was followed by 1-minute conditioning sessions where mice alternated between interactions with a CD-1 aggressor, present in one of the chambers (CS+), and an empty chamber (CS-, **Fig. 1C**), as described in detail in the materials and methods section. Overall, mice showed no initial preference for either chamber during the Pre-Test (**Fig. 1D**). Following two conditioning sessions, mice still showed no preference for the CS+ or CS− chamber (Avoidance Test 1). However, after four conditioning sessions (Avoidance Test 2), mice spent significantly less time in the CS+ than in the CS− chamber, indicating the development of avoidance toward the CD-1-paired context (**Fig. 1D**). Despite this group-level effect, we observed substantial inter-individual variability, with 2 out of 10 mice showing a preference, rather than avoidance, for the CD-1-paired chamber. Three days later, mice were tested again in the empty arena to assess long-term memory and showed a trend toward avoidance of the CS+ chamber, suggesting partial retention of the conditioned place aversion (**Fig. 1D**).

We wondered whether reduced avoidance learning in these specific individuals could be due to low levels of aggression during conditioning sessions. To answer this question, defensive and offensive behaviors, displayed by BL6 and CD-1 mice respectively, were manually scored. CD-1 mice displayed a rich repertoire of offensive behaviors, with attacks, biting and aggressive sniffing towards BL6 mice (**Fig. 1E-F** and **S1A-D**), whereas BL6 mice showed an array of defensive behaviors including freezing, defensive behavior and escaping (**Fig. 1E-F** and **S1E-H**). Indeed, the amount of attack shown by CD-1 mice and the amount of defensive behavior displayed by BL6 mice positively correlated, suggesting complementary responding within the dyad (**Fig. S1I**). Generally, offensive and defensive behaviors did not habituate over the 4 conditioning days, indicating temporal stability of the encounters (**Fig. S1A-H** for individual behaviors and **Fig. 1F** for summary). Correlation analysis between the amount of defensive and offensive behaviors displayed during the conditioning sessions and the Preference Index (PI) measured in Avoidance Test 2 revealed no significant association between these variables (**Fig. 1G**). These findings indicate that the extent of social conditioned place aversion is not directly related to the level of aggression experienced during conditioning in the sCPA.

### Social aversive conditioning leads to persistent behavioral changes and generalized avoidance for unfamiliar CD-1 mice

To determine whether avoidance learning was driven specifically by the aversive social interaction rather than by repeated exposure to experimental manipulations and handling procedures, we tested an independent cohort of mice in which the CD-1 aggressor was replaced by an object during conditioning (**Fig. 2A-B**). For this second experiment, mice were additionally subjected to a battery of behavioral assessments before and after the conditioning procedure, allowing us to characterize broader behavioral changes associated with the sCPA (**Fig. S2A**). As in the original cohort, mice exposed to CD-1 aggressors developed avoidance following four conditioning sessions, reproducing the behavioral phenotype observed in our initial experiments (**Fig. 2C-D**). In contrast, object-conditioned mice showed no chamber preference during the Pre-Test or after two conditioning sessions (Avoidance Test 1), and failed to develop avoidance of the CS+ chamber after four conditioning sessions (Avoidance Test 2) or during the Long-term memory Test (**Fig. 2B**). Together, these findings demonstrate that contextual avoidance in the sCPA task depends on the aversive social experience and cannot be explained by repeated exposure to a non-social stimulus.

**Figure 2 -.**
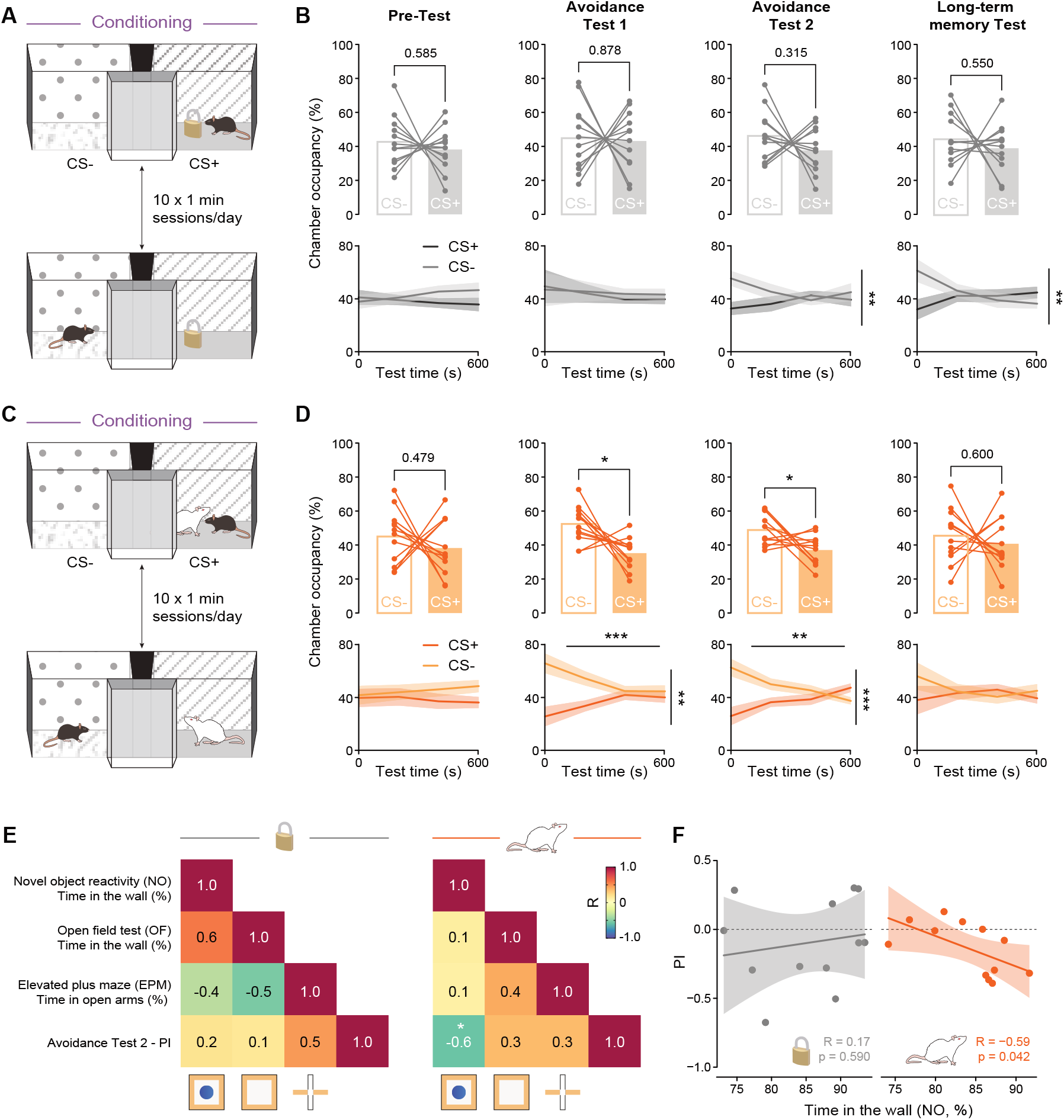
Social aversive conditioning drives place avoidance and can be linked to individual variability in anxiety-related variables. **A**, Control conditioning procedure. Mice were conditioned by pairing one chamber (CS+) with an inanimate object, while the opposite chamber (CS−) remained empty. Conditioning was performed in alternating sessions (10 ✕ 1-min sessions per day). **B**, Behavioral performance following object-based conditioning (control). Top: occupancy of the CS− and CS+ chambers during the Pre-test, Avoidance Test 1, Avoidance Test 2, and Long-term memory Test. Each line represents an individual animal. Pre-Test (paired t-test): t(11) = 0.563, p = 0.585; Avoidance Test 1 (paired t-test): t(11) = 0.158, p = 0.878; Avoidance Test 2 (paired t-test): t(11) = 1.053, p = 0.315; Long-term memory Test (paired t-test): t(11) = 0.617, p = 0.550. Bottom: temporal occupancy dynamics across the test sessions. Shaded areas represent mean ± SEM. Pre-Test (RM two-way ANOVA with Greenhouse-Geisser correction): chamber effect, F(1,22) = 0.625, p = 0.437; time effect, F(1.706, 37.532) = 0.159, p = 0.820; chamber ✕ time interaction, F(1.706, 37.532) = 2.267, p = 0.125. Avoidance Test 1 (RM two-way ANOVA with Greenhouse-Geisser correction): chamber effect, F(1, 22) = 0.049, p = 0.827; time effect, F(1.423, 31.310) = 0.480, p = 0.559; chamber ✕ time interaction, F(1.423, 31.310) = 0.106, p = 0.832. Avoidance Test 2 (RM two-way ANOVA): chamber effect, F(1, 22) = 2.164, p = 0.155; time effect, F(3, 66) = 0.342, p = 0.795; chamber ✕ time interaction, F(3, 66) = 4.143, p = 0.009. Long-term memory Test (RM two-way ANOVA with Greenhouse-Geisser correction): chamber effect, F(1, 22) = 0.747, p = 0.397; time effect, F(1.770, 38.933) = 0.940, p = 0.389; chamber ✕ time interaction, F(1.770, 38.933) = 7.360, p = 0.003. **C**, Social Conditioned Place Aversion procedure. Mice were conditioned by pairing one chamber (CS+) with an aggressive CD-1 mouse and the opposite chamber (CS−) with no social stimulus. Conditioning was performed in alternating sessions (10 ✕ 1-min sessions per day). **D**, Behavioral performance following aversive social conditioning (sCPA). Top: occupancy of the CS− and CS+ chambers during the Pre-test, Avoidance Test 1, Avoidance Test 2, and Long-term memory Test. Each line represents an individual animal. Pre-Test (paired t-test): t(11) = 0.733, p = 0.479; Avoidance Test 1 (paired t-test): t(11) = 3.028, p = 0.011; Avoidance Test 2 (paired t-test): t(11) = 2.474, p = 0.031; Long-term memory Test (paired t-test): t(11) = 0.539, p = 0.600. Bottom: temporal occupancy dynamics across the test sessions. Shaded areas represent mean ± SEM. Pre-Test (RM two-way ANOVA with Greenhouse-Geisser correction): chamber effect, F(1, 22) = 1.050, p = 0.317; time effect, F(1.903, 41.872) = 0.068, p = 0.927; chamber ✕ time interaction, F(1.903, 41.872) = 0.685, p = 0.503. Avoidance Test 1 (RM two-way ANOVA with Greenhouse-Geisser correction): chamber effect, F(1, 22) = 17.412, p < 0.001; time effect, F(2.026, 44.579) = 0.156, p = 0.859; chamber ✕ time interaction, F(2.026, 44.579) = 5.707, p = 0.006. Avoidance Test 2 (RM two-way ANOVA with Greenhouse-Geisser correction): chamber effect, F(1, 22) = 10.687, p = 0.004; time effect, F(1.797, 39.537) = 0.101, p = 0.885; chamber ✕ time interaction, F(1.797, 39.537) = 9.077, p < 0.001. Long-term memory Test (RM two-way ANOVA with Greenhouse-Geisser correction): chamber effect, F(1, 22) = 0.559, p = 0.463; time effect, F(1.863, 40.979) = 0.258, p = 0.758; chamber ✕ time interaction, F(1.863, 40.979) = 1.451, p = 0.246. **E**, Correlation matrices showing relationships between anxiety-like behavior measured in the reactivity to a Novel Object (NO), Open Field (OF), and Elevated Plus Maze (EPM) tests, and avoidance performance following object-based (left) or aversive social (right) conditioning. Values indicate Pearson correlation coefficients. PI, Preference Index calculated as (“CS+” − “CS−”)/(“CS+” + “CS−”). **F**, Relationship between anxiety-like behavior, quantified as time spent in the wall zone during the Novel Object test, and avoidance/preference for the chambers as assessed by the Preference Index (PI). Left: object-conditioned animals. Right: socially conditioned animals. Each point represents one animal. Statistical test: Pearson correlation. Object conditioning: R = 0.17, p = 0.590; aversive social conditioning: R = −0.59, p = 0.042.

**Figure 3 -.**
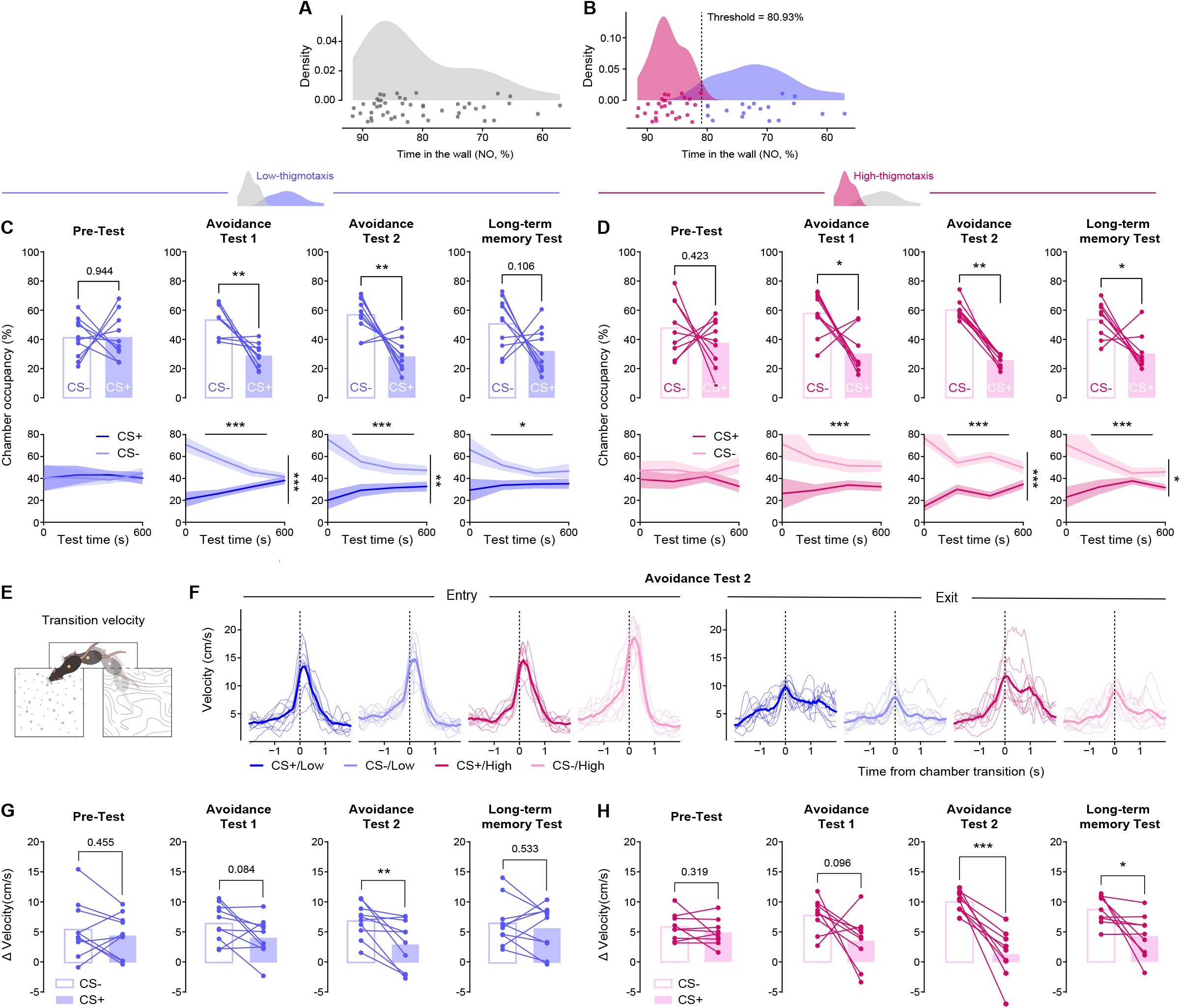
Baseline novelty-induced thigmotaxis identifies mice with distinct susceptibility to social avoidance learning. **A**, Distribution of anxiety-like behavior measured as time spent in the wall zone (*i.e.* thigmotaxis) during the Novel Object (NO) test across all animals. **B**, Classification of animals into low-and high-thigmotaxis groups using a two-component Gaussian mixture model fitted to the distribution of wall occupancy during the NO test. The classification threshold was defined as the intersection between the two fitted distributions (threshold = 80.93%). **C**, Behavioral performance of low-thigmotaxis animals during the social Conditioned Place Aversion task. Top: occupancy of the CS− and CS+ chambers during the Pre-Test, Avoidance Test 1, Avoidance Test 2, and Long-term memory Test. Each line represents an individual animal. Pre-Test (paired t-test): t(9) = −0.072, p = 0.944; Avoidance Test 1 (paired t-test): t(9) = 4.314, p = 0.002; Avoidance Test 2 (paired t-test): t(9) = 3.975, p = 0.003; Long-term memory Test (paired t-test): t(9) = 1.797, p = 0.106. Bottom: temporal occupancy dynamics across the test sessions. Shaded areas represent mean ± SEM. Pre-Test (RM two-way ANOVA with Greenhouse-Geisser correction): chamber effect, F(1, 18) = 0.009, p = 0.924; time effect, F(1.522, 27.393) = 0.032, p = 0.937; chamber ✕ time interaction, F(1.522, 27.393) = 0.135, p = 0.818. Avoidance Test 1 (RM two-way ANOVA with Greenhouse-Geisser correction): chamber effect, F(1, 18) = 36.094, p < 0.001; time effect, F(1.706, 30.708) = 0.968, p = 0.379; chamber ✕ time interaction, F(1.706, 30.708) = 10.961, p < 0.001. Avoidance Test 2 (RM two-way ANOVA): chamber effect, F(1, 18) = 31.066, p < 0.001; time effect, F(3, 54) = 0.865, p = 0.465; chamber ✕ time interaction, F(3, 54) = 5.572, p = 0.002. Long-term memory Test (RM two-way ANOVA with Greenhouse-Geisser correction): chamber effect, F(1, 18) = 6.235, p = 0.022; time effect, F(1.394, 25.092) = 1.135, p = 0.318; chamber ✕ time interaction, F(1.394, 25.092) = 3.501, p = 0.060. **D**, Behavioral performance of high-thigmotaxis animals during the social Conditioned Place Aversion task. Top: occupancy of the CS− and CS+ chambers during the Pre-test, Avoidance Test 1, Avoidance Test 2, and Long-term memory Test. Each line represents an individual animal. Pre-Test (paired t-test): t(8) = 0.845, p = 0.423; Avoidance Test 1 (Wilcoxon test): W = −37.000, p = 0.027; Avoidance Test 2 (Wilcoxon test): W = −45.000, p = 0.004; Long-term memory Test (Wilcoxon test): W = −37.000, p = 0.027. Bottom: temporal occupancy dynamics across the test sessions. Shaded areas represent mean ± SEM. Pre-Test (RM two-way ANOVA): chamber effect, F(1, 16) = 1.398, p = 0.254; time effect, F(3, 48) = 0.013, p = 0.998; chamber ✕ time interaction, F(3, 48) = 1.481, p = 0.231. Avoidance Test 1 (two-way RM ANOVA with Greenhouse-Geisser correction): chamber effect, F(1, 16) = 15.442, p = 0.001; time effect, F(1.491, 23.854) = 0.349, p = 0.647; chamber ✕ time interaction, F(1.491, 23.854) = 1.409, p = 0.259. Avoidance Test 2 (RM two-way ANOVA with Greenhouse-Geisser correction): chamber effect, F(1, 16) = 173.725, p < 0.001; time effect, F(2.122, 33.947) = 0.269, p = 0.779; chamber ✕ time interaction, F(2.122, 33.947) = 8.443, p < 0.001. Long-term memory Test (RM two-way ANOVA with Greenhouse-Geisser correction): chamber effect, F(1, 16) = 15.577, p = 0.001; time effect, F(1.504, 24.066) = 0.767, p = 0.441; chamber ✕ time interaction, F(1.504, 24.066) = 4.706, p = 0.027. **E**, Schematic illustrating transition velocity, defined as the instantaneous locomotor speed surrounding transitions between compartments. **F**, Velocity profiles aligned to chamber transitions during Avoidance Test 2 in low-and high-thigmotaxis mice. Traces show movement velocity around entrance into (left) and exit from (right) the CS+ and CS− chambers. Time 0 (dashed line) indicates the chamber transition. Thin lines represent individual mice and thick lines represent the group average. **G**, Change in transition velocity (ΔVelocity) during testing in low-thigmotaxis animals. Each line represents an individual animal. Pre-Test (paired t-test): t(9) = 0.781, p = 0.455; Avoidance Test 1 (paired t-test): t(9) = 1.941, p = 0.084; Avoidance Test 2 (paired t-test): t(9) = 3.380, p = 0.008; Long-term memory Test (paired t-test): t(9) = 0.649, p = 0.533. **H**, Change in transition velocity (ΔVelocity) during testing in high-thigmotaxis animals. Each line represents an individual animal. Pre-Test (paired t-test): t(8) = 1.063, p = 0.319; Avoidance Test 1 (paired t-test): t(8) = 1.888, p = 0.096; Avoidance Test 2 (paired t-test): t(8) = 5.683, p < 0.001; Long-term memory Test (paired t-test): t(8) = 2.663, p = 0.029.

We next took advantage of the longitudinal behavioral characterization of this cohort to determine whether repeated social aversive conditioning induced behavioral changes beyond avoidance of the conditioned context. Following the procedure, both control and sCPA mice showed reduced locomotor activity in the Open Field (OF, **Fig. S2B, S2C** & **S2D** left panels). However, only socially conditioned mice showed a significant reduction in center zone exploration, consistent with a selective change in anxiety-related exploratory behavior (**Fig. S2C** & **S2D** middle panels). Reactivity to a Novel Object was also assessed before and after conditioning. While object-conditioned controls increased their exploration of the center containing the novel object, this change was not observed following sCPA (NO, **Fig. S2E, S2F** & **S2G** middle panels). In the Novel Object Recognition test, control mice preferentially explored the novel object, whereas this discrimination was weaker following sCPA, suggesting a modest alteration in recognition memory (NOR, **Fig. S2H–J**). To determine whether the social experience produced consequences that persisted beyond contextual avoidance, mice were exposed to a CD-1 mouse one month after conditioning (**Fig. S2K–N**). Socially conditioned mice displayed reduced interaction with the CD-1 compared with control animals, indicating that the consequences of the aversive social experience persisted well beyond the conditioning period and suggesting a generalized fear towards unfamiliar CD-1 aggressors (**Fig. S2N**). Together, these findings show that the sCPA procedure not only consistently induces conditioned place aversion but is also accompanied by longer-lasting alterations across anxiety-related, cognitive, and social behavioral domains.

We next sought to better characterize the behavioral expression of conditioned responses in the sCPA. Because freezing is a widely established readout of conditioned fear, we quantified freezing within each chamber across the different test phases of the sCPA procedure (**Fig. S3A**). Object-conditioned mice showed no chamber-specific differences in freezing at any test session (**Fig. S3B-C**). In contrast, CD-1-conditioned mice displayed significant differences in freezing during both Avoidance Tests 1 and 2, but not during the Pre-Test or Long-term memory Test (**Fig. S3B** and **D**). Surprisingly, freezing was higher in the CS− than in the CS+ chamber. To further investigate this finding, we asked whether conditioning instead altered locomotor behavior within the two contexts. Object-conditioned mice showed no differences in mean moving velocity between the CS+ and CS− chambers across test sessions (**Fig. S3E**). CD-1-conditioned mice, however, showed a chamber-dependent difference in moving velocity during Avoidance Test 1, with higher velocity in the CS+ chamber (**Fig. S3F**). Additional behavioral readout comparisons between chambers for both groups, and specifically regarding pauses, sniffing, grooming and rearing, are reported in **Supplementary Table 4**. Together, these findings suggest that the behavioral response elicited by the socially conditioned context is not merely dominated by alterations in freezing but instead entails changes in mice’s exploratory strategies.

### Novelty-induced thigmotaxis predicts individual differences in avoidance learning

The marked inter-individual variability in avoidance observed across mice that was not associated with the amount of aggression received during conditioning, prompted us to ask whether avoidance learning in the sCPA could be related to behavioral traits present before conditioning. Taking advantage of the pre-conditioning behavioral characterization of this cohort, we examined anxiety-related measures obtained from the OF, NO and elevated plus maze (EPM) tests (**Fig. S2A**). Correlation analysis revealed little correspondence among most behavioral measures, indicating that these assays captured distinct aspects of emotional behavior (**Fig. 2E**). Notably, neither thigmotaxis (*i.e*. when mice avoid open areas and prefer to stay close to the walls) in the OF nor time spent in the open arms of the EPM correlated with avoidance learning in neither object-conditioned nor CD-1-conditioned mice. In contrast, thigmotaxic behavior during the NO test showed a significant negative correlation with the PI measured in the CD-1-conditioned group (**Fig. 2E-F**). This NO readout consists of a novelty aspect, *i.e.* novel object that the mice are curious to explore, but placed in an anxiogenic part of the arena, *i.e.* the center of an open field. Mice that spent more time exploring the wall zone during exposure to the novel object subsequently displayed stronger avoidance of the CD-1-paired chamber. These findings identify anxiety-related readouts and namely novelty-induced thigmotaxis, as potential predictors of susceptibility to social avoidance learning.

### Baseline novelty-induced thigmotaxis identifies mice with distinct susceptibility to social avoidance learning

To challenge this relationship further, we designed an experiment with a larger cohort of mice that were exposed to anxiety-like behavioral assessments. Subsequently, we examined the distribution of time spent in the wall zone during the NO test. Density analysis revealed a bimodal distribution, allowing the identification of two behavioral subpopulations separated by a threshold of 80.93% wall occupancy (**Fig. 3A-B**). Based on this threshold, mice were prospectively classified into low-and high-thigmotaxis groups (*i.e.* low and high anxiety-like behavior respectively) before undergoing the sCPA. Both groups showed no preference for either chamber during the Pre-Test (**Fig. 3C-D**). Following conditioning, however, clear differences emerged. Although both groups exhibited conditioned place aversion already during Avoidance Test 1, only mice with high baseline thigmotaxis displayed persistent avoidance of the CS+ chamber during the Long-term memory Test (**Fig. 3C**-**D**).

Because freezing and mean moving velocity within each chamber were shown previously to be distinctly modulated in sCPA vs. control mice (*i.e.* object-conditioned), we then asked whether locomotor parameters would show group-specific expression patterns in low-and high-thigmotaxis mice. Mean moving velocity was increased in the CS+ chamber during the Avoidance Test 2 in both groups (**Fig. S4A-B**), similarly to the Avoidance Test 1 in our previous sCPA cohort (**Fig. S3F**). We next further investigated velocity signatures pertaining to chamber transitions (**Fig. S4C-F**). Both high-and low-thigmotaxis mice showed slower entrance speed and more rapid exits from the CS+ chamber as compared to the CS-context during Avoidance Test 2 (**Fig. S4C-F**), but interestingly only the high-thigmotaxis group entered the CS+ chamber slower during the Long-term memory Test (**Fig. S4D**). We thus calculated the change in movement velocity between chamber entry and exit (ΔVelocity) (**Fig. 3E-F**). Whereas both low-and high-thigmotaxis mice exhibited a marked reduction in ΔVelocity in the CS+ chamber during Avoidance Test 2, only high-thigmotaxis mice maintained this effect during the Long-term memory Test (**Fig. 3G-H**). Thus, in addition to developing persistent, conditioned avoidance, high-thigmotaxis mice also displayed a more pronounced change in locomotor behavior when navigating the aversively conditioned context. However, when re-exposed to a novel CD-1 mouse, high-thigmotaxis mice behaved similarly to low-thigmotaxis mice showing generalized avoidance as previously reported (**Fig. S4G-J**).

Together, these findings indicate that prior high anxiety-related traits are associated with specific alterations in contextual social fear processing, denoted by persistent long-term memory in the high-thigmotaxis group. Moreover, they corroborate data from the previous cohort where distinct behavioral locomotor strategies are adopted when interacting with the aversively conditioned environment.

### Pre-test behavioral features predict post-test avoidance outcomes

Having identified novelty-induced thigmotaxis as a predictor of social avoidance learning, we next asked whether susceptibility to social avoidance could be predicted from the broader behavioral state expressed before conditioning. To test this, we trained a gradient-boosted decision tree model (XGBoost) using exclusively Pre-Test variables capturing ethological states and locomotor dynamics across the apparatus and within the corridor, while excluding chamber-specific information related to conditioned or non-conditioned contexts (**Fig. 4A**). The target variable was the PI, reflecting the relative avoidance of the aggressor-paired chamber. The model predicted inter-individual variability in avoidance behavior above chance levels (mean cross-validated Pearson’s R = 0.45, permutation p = 0.04, **Fig. 4B**). Although predictive accuracy was modest, these findings indicate that stable individual differences in behavioral state prior to conditioning contribute to variability in social aversive learning outcomes. Interestingly, the model could not efficiently predict post-test behavior in the control procedure where animals were exposed to an object in one of the chambers instead of a CD-1 mouse (**Fig. 4C**), suggesting that the predictive relationship identified here is specific to the social stress-based CPA paradigm and does not extend to more neutral contexts. Analysis of feature importance revealed that the model relies primarily on a small subset of behavioral variables (**Fig. 4D**). Notably, measures related to rearing and sniffing behavior, locomotor activity, and corridor occupancy during the final phase of the session (450–600 s) were among the top predictors. To further interpret how these features influence model predictions, we examined their relationship with the predicted outcome (**Fig. 4E**). Higher levels of rearing and locomotor activity were associated with reduced avoidance (higher PI), whereas lower activity levels predicted stronger avoidance. Remarkably, behavioral traits such as rearing remained highly stable across testing phases (**Fig. 4F**), suggesting that they reflect trait-like characteristics of individual animals.

**Figure 4 -.**
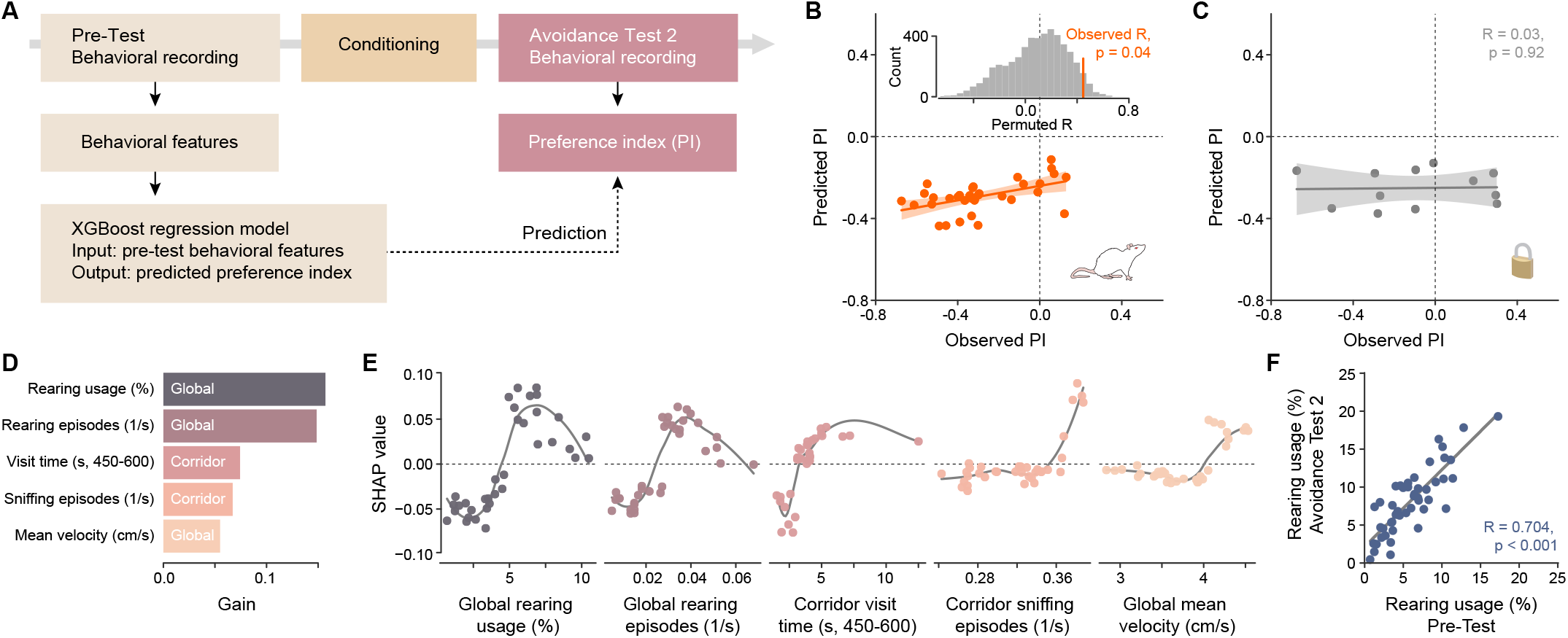
Pre-Test behavioral features predict susceptibility to social aversive learning. **A**, Schematic of the machine-learning pipeline. Behavioral features were extracted from the Pre-Test session and used to train a gradient-boosted decision tree model (XGBoost) to predict the Preference Index (PI) measured during Avoidance Test 2. Only behavioral variables capturing ethological states and locomotor dynamics across the apparatus and corridor were included (see **Materials and Methods** for more information). **B**, Representative out-of-fold (OOF) prediction showing the relationship between observed and predicted PI values. The inset shows the null distribution obtained from 5,000 target-label permutations. The horizontal orange line indicates the observed mean cross-validated Pearson correlation coefficient (R = 0.45, permutation p = 0.04). The representative OOF run shown yielded Pearson’s R = 0.51 and Spearman’s ρ = 0.51. **C**, Application of the same modeling approach to the control procedure in which animals were exposed to an object instead of a CD-1 aggressor mouse during conditioning. No significant predictive relationship was observed. **D**, Relative feature importance (gain) of the top 5 behavioral predictors identified by the model. **E**, SHAP (SHapley Additive exPlanations) dependence plots illustrating the contribution of the top predictive features to model output. Positive SHAP values indicate increased predicted PI (reduced avoidance), whereas negative SHAP values indicate decreased predicted PI (stronger avoidance). **F**, Relationship between rearing usage measured during the Pre-Test and Avoidance Test 2 (Pearson’s R = 0.704, p < 0.001).

We next examined whether the main predictive features were related to novelty-induced thigmotaxis or represented largely independent behavioral dimensions. Thigmotaxis tended to correlate negatively with rearing measures, although these associations were modest and did not reach statistical significance (**Fig. S5**). These data denote that the association between anxiety-like behavior and other locomotor/exploration features is not straightforward; rather, social avoidance may depend upon a multifactorial behavioral profile contingent on specific encoding, conditioning and memory testing stages during contextual social fear processing.

Taken together, these results demonstrate that individual differences in contextual social fear learning can be shaped with the influence of pre-existing behavioral metrics and exploratory/locomotor dynamics.

### VTA*^Vgat^* cells are essential for the acquisition of social stress-induced conditioned place aversion

Having established the sCPA as a robust paradigm for social stress context processing, we next sought to characterize the brain regions engaged during the task. To this end, we quantified the expression of the immediate early gene c-Fos in selected brain areas implicated in contextual and aversive learning, including the dorsal hippocampal formation^50,51^ the lateral and basolateral amygdala,^52,53^ and the ventral tegmental area (VTA).^54^ Compared with object-conditioned control mice, c-Fos density showed region-specific changes in the sCPA condition following the Avoidance Test 1. No significant changes were detected in the lateral or basolateral amygdala (**Fig. 5A**). Although several hippocampal regions, such as the dentate gyrus (DG) and CA1, as well as the hippocampus as a whole (HPC), showed increased c-Fos density, the VTA was the only region showing a reduction in c-Fos density (**Fig. 5A**). This distinct decrease prompted us to investigate further the implicated VTA neuronal subpopulations. For this purpose, we performed double immunostaining for tyrosine hydroxylase (TH) and c-Fos. The decrease in overall VTA c-Fos expression was selectively driven by a reduction in c-Fos-positive/TH-positive (c-Fos^+^/TH^+^) neurons, while the density of c-Fos-positive/TH-negative (c-Fos^+^/TH^-^) neurons remained stable across conditions (**Fig. 5B-C**). These findings indicate that dopaminergic, but not TH^-^, VTA neurons undergo changes in activity during social conditioned place aversion processing. Indeed, both direct and indirect (via VTA GABA activation) VTA DA neuron inhibition were shown previously to lead to conditioned place aversion.^37^ Prior important work has also demonstrated that VTA GABAergic inhibition leads to disinhibition of VTA DA neurons.^55–57^ Thus, we sought to selectively inhibit the VTA GABAergic population and assess whether this would perturb sCPA processing. To this end, we expressed hM4Di selectively in VTA GABA neurons of Vgat-cre mice, reasoning that transient inhibition of these interneurons during conditioning would interfere with learning of the task. Control Vgat-cre mice received a Cre-dependent mCherry virus to control for stereotaxic surgery and potential secondary effects of systemic Clozapine-N-oxide (CNO). Following at least three weeks of viral expression, mice underwent the sCPA protocol and received CNO 30 minutes before each conditioning session (**Fig. 5D**).

**Figure 5 -.**
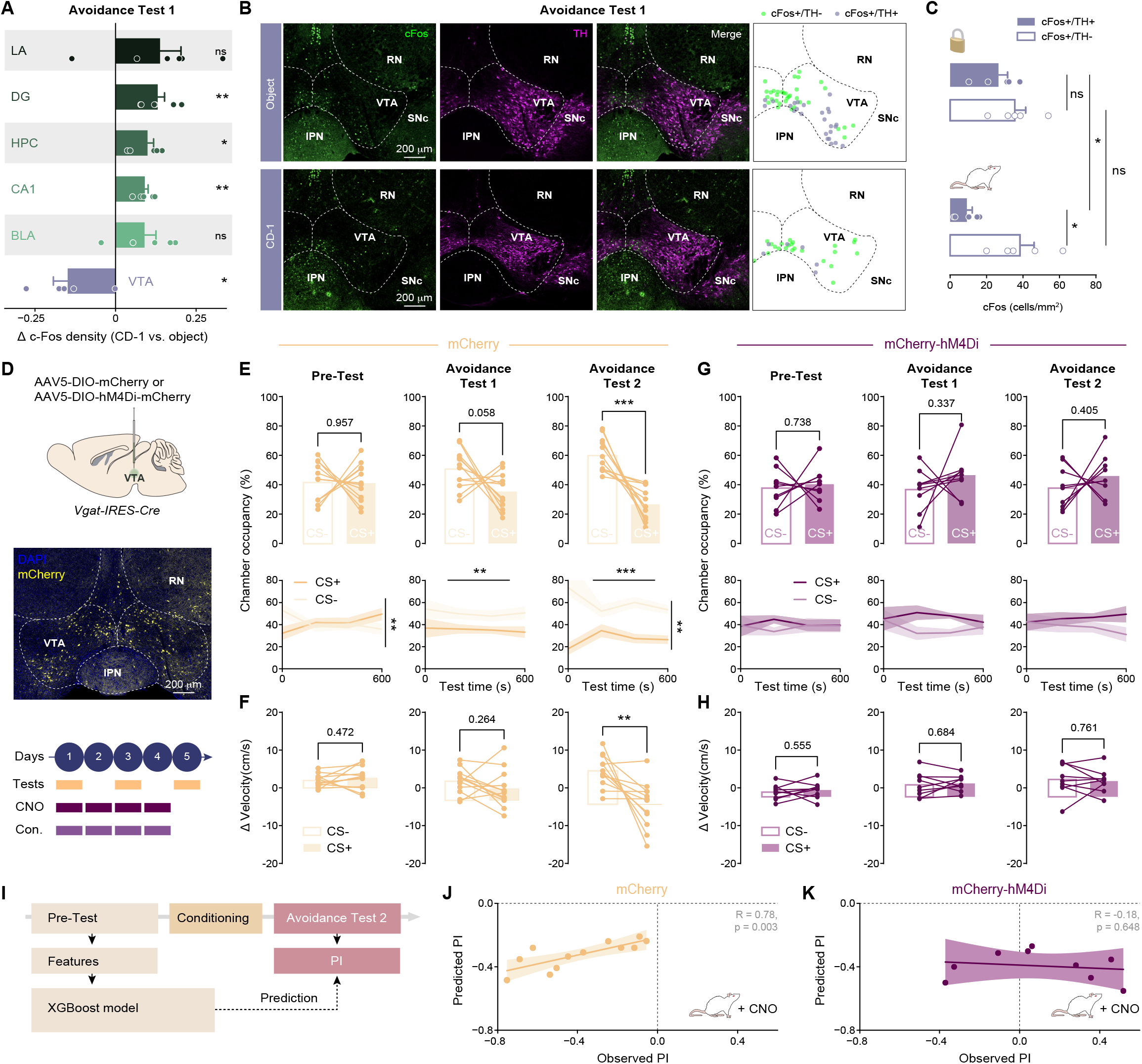
Inhibition of VTA^Vgat^ neurons during conditioning impairs the acquisition of social conditioned place aversion. **A,** Changes in c-Fos^+^ cell density in sCPA animals across selected brain regions following the Avoidance Test 1, expressed relative to the object-conditioned group. Positive and negative values indicate increased and decreased c-Fos density relative to controls, respectively. LA, lateral amygdala; DG, dentate gyrus; HPC, hippocampus; CA1, cornu ammonis 1; BLA, basolateral amygdala; VTA, ventral tegmental area. One-sample t-test against 0, Benjamini– Hochberg False Discovery Rate (FDR)-corrected across regions (LA: p = 0.087, DG: p = 0.005, HPC: p = 0.006, CA1: p = 0.002, BLA: p = 0.060, VTA: p = 0.043). **B**, Representative images of c-Fos (green) and tyrosine hydroxylase (TH; magenta) immunostaining in the VTA of control (top) and sCPA (bottom) groups, with corresponding merged images and spatial distribution of c-Fos+/TH− and c-Fos+/TH+ cells. Dashed lines delineate anatomical regions. RN, red nucleus; SNc, substantia nigra pars compacta; IPN, interpeduncular nucleus. Scale bar, 200 μm. **C**, Quantification of c-Fos+/TH+ and c-Fos+/TH− cell density in the VTA following the Avoidance Test 1 for the sCPA and control group. Statistical test: RM two-way ANOVA with Greenhouse-Geisser correction followed by Sidak post hoc test. Group effect, F(1,8) = 6.358, p = 0.036; cell type effect, F(1.000, 8.000) = 7.938, p = 0.023; group ✕ cell type interaction, F(1.000, 8.000) = 2.113, p = 0.184. **D**, Experimental design and timeline. Vgat-IRES-Cre (Vgat-cre) mice received bilateral injections of AAV5-hSyn-DIO-mCherry or AAV5-hSyn-DIO-hM4Di-mCherry into the VTA. Mice underwent the social Conditioned Place Aversion paradigm while receiving CNO before each conditioning session. Behavioral performance was assessed during the Pre-Test, Avoidance Test 1 and Avoidance Test 2. **E**, Behavioral performance of control (mCherry) mice. Top: occupancy of the CS− and CS+ chambers during the Pre-test, Avoidance Test 1 and Avoidance Test 2. Each line represents an individual animal. Pre-Test (paired t-test): t(11) = 0.055, p = 0.957; Avoidance Test 1 (paired t-test): t(11) = 2.114, p = 0.058; Avoidance Test 2 (paired t-test): t(11) = 5.316, p < 0.001. Bottom: temporal occupancy dynamics across the test sessions. Shaded areas represent mean ± SEM. Pre-Test (RM two-way ANOVA with Greenhouse-Geisser correction): chamber effect, F(1, 22) = 0.008, p = 0.931; time effect, F(2.167, 47.667) = 0.240, p = 0.805; chamber ✕ time interaction, F(2.167, 47.667) = 6.740, p = 0.002. Avoidance Test 1 (RM two-way ANOVA with Greenhouse-Geisser correction): chamber effect, F(1, 22) = 8.538, p = 0.008; time effect, F(2.004, 44.087) = 0.232, p = 0.794; chamber ✕ time interaction, F(2.004, 44.087) = 0.101, p = 0.905. Avoidance Test 2 (RM two-way ANOVA with Greenhouse-Geisser correction): chamber effect, F(1, 22) = 53.154, p < 0.001; time effect, F(2.024, 44.536) = 0.550, p = 0.583; chamber ✕ time interaction, F(2.024, 44.536) = 5.905, p = 0.005. **F**, Change in transition velocity (ΔVelocity) in control (mCherry) mice. Each line represents an individual animal. Pre-Test (paired t-test): t(11) = −0.745, p = 0.472; Avoidance Test 1 (paired t-test): t(11) = 1.177, p = 0.264; Avoidance Test 2 (paired t-test): t(11) = 3.517, p = 0.005. **G**, Behavioral performance of hM4Di-expressing mice during social Conditioned Place Aversion. Top: occupancy of the CS− and CS+ chambers during the Pre-Test, Avoidance Test 1 and Avoidance Test 2. Each line represents an individual animal. Pre-Test (paired t-test): t(8) = −0.346, p = 0.738; Avoidance Test 1 (paired t-test): t(8) = −1.021, p = 0.337; Avoidance Test 2 (paired t-test): t(8) = −0.880, p = 0.405. Bottom: temporal occupancy dynamics across the test sessions. Shaded areas represent mean ± SEM. Pre-Test (RM two-way ANOVA): chamber effect, F(1, 16) = 0.213, p = 0.651; time effect, F(3, 48) = 0.002, p = 1.000; chamber ✕ time interaction, F(3, 48) =0.503, p = 0.682. Avoidance Test 1 (RM two-way ANOVA with Greenhouse-Geisser correction): chamber effect, F(1, 16) = 2.002, p = 0.176; time effect, F(1.986, 31.778) = 0.319, p = 0.728; chamber ✕ time interaction, F(1.986, 31.778) = 1.186, p = 0.318. Avoidance Test 2 (RM two-way ANOVA): chamber effect, F(1, 16) = 1.486, p = 0.241; time effect, F(3, 48) = 0.088, p = 0.966; chamber ✕ time interaction, F(3, 48) = 1.253, p = 0.301. **H**, Change in transition velocity (ΔVelocity) in hM4Di-expressing mice. Each line represents an individual animal. Pre-Test (paired t-test): t(8) = −0.616, p = 0.555; Avoidance Test 1 (paired t-test): t(8) = −0.422, p = 0.684; Avoidance Test 2 (paired t-test): t(8) = 0.314, p = 0.761. **I**, Schematic of the prediction analysis. The XGBoost model previously trained on Pre-Test behavioral features (Fig. 4) was applied to predict the Preference Index (PI) measured during Avoidance Test 2 in control and hM4Di mice. Neither cohort was used for model training. **J**, Relationship between observed and model-predicted PI in control mCherry mice (Pearson’s R = 0.78, p = 0.003). **K**, Relationship between observed and model-predicted PI in hM4Di mice (Pearson’s R = −0.18, p = 0.648).

Control virus-injected mice reproduced the behavioral phenotype observed in previous cohorts, showing no chamber preference during the Pre-Test and robust avoidance of the CS+ chamber during Avoidance Tests (**Fig. 5E**). In addition, control mCherry mice displayed the previously observed consistent reduction in transition velocity toward the CS+ chamber during Avoidance Test 2 (**Fig. 5F**). These findings confirm that the sCPA paradigm is robust across mouse lines and is not confounded by stereotaxic surgery or systemic CNO administration. In contrast, mice expressing hM4Di in VTA GABA neurons failed to develop conditioned place avoidance. These animals showed no preference for the CS− over the CS+ chamber during either Avoidance Tests (**Fig. 5G**). Likewise, the reduction in transition velocity typically associated with intact avoidance learning was absent following chemogenetic inhibition of VTA GABA neurons (**Fig. 5H**), indicating that VTA GABAergic activity during conditioning is required for the context processing in the sCPA.

We then tested whether inhibiting VTA GABA neurons during conditioning altered the relationship between pre-existing behavioral state and subsequent avoidance learning. To this end, we applied the behavioral prediction model, trained exclusively on Pre-Test behavioral features from previous cohorts (see **Fig. 4**), to both control and hM4Di mice (**Fig. 5I**). In control animals, the model accurately predicted post-test avoidance, confirming its ability to generalize across independent cohorts (**Fig. 5J**). In contrast, predictive performance was markedly reduced in hM4Di mice (**Fig. 5K**). Thus, inhibition of VTA GABA neurons during conditioning disrupted the relationship between baseline behavioral features and later avoidance outcome. These findings indicate that our model, using pre-existing individual traits, predicts behaviorally meaningful learning outcomes with accuracy and specificity, rather than capturing random variation or impaired contextual processing readouts resulting from chemogenetic manipulation.

Because inhibition of VTA GABA neurons can increase locomotor activity,^58^ we used a relatively low dose of CNO (0.3 mg/kg). However, we performed additional experiments to assess potential behavioral consequences of the manipulation. When tested 30 minutes after CNO administration in a three-chamber assay (**Fig. S6A**), hM4Di mice displayed increased locomotor activity compared with control mice during both habituation and social preference testing (**Fig. S6B**), confirming the expected locomotor effect of the manipulation. Despite this increase, both groups preferentially explored the enclosure containing a juvenile mouse over the empty enclosure and showed a juvenile exploration ratio above chance, indicating that chemogenetic inhibition did not impair general social preference (**Fig. S6C-D**). Importantly, no differences in overall distance traveled were detected between control and hM4Di mice during the sCPA test sessions, which were performed without acute CNO administration (**Fig. S6E-F**). Thus, the absence of conditioned avoidance in hM4Di mice is unlikely to reflect a persistent alteration in locomotor activity during testing.

Finally, we examined whether VTA GABA inhibition influenced the physiological response to an acute social defeat. Mice received CNO 30 minutes before a 5-minutes encounter with a novel CD-1 aggressor, and circulating corticosterone was measured 30 minutes after the encounter. hM4Di mice displayed lower corticosterone levels than control mice (**Fig. S6G**), indicating that inhibition of VTA GABA neurons attenuates the endocrine stress response to social defeat.

## Discussion

The work presented here establishes and characterizes a protocol of conditioned place aversion based on social stress where mice show avoidance for a chamber paired with a CD-1 aggressor (CS+) during several conditioning sessions, notably with inter-individual variability. Investigating the amount of aggression received during conditioning sessions revealed no significant relationship of offensive/defensive behaviors with avoidance learning, indicating the influence of other underlying factors. Indeed, prior anxiety-like behavior was associated with avoidance learning, with highly anxious individuals (high thigmotaxis group) showing persistent avoidance for the chamber previously paired with a CD-1 mouse. Expanding on our question regarding prior individual traits in shaping avoidance learning in the sCPA, we next investigated whether Pre-Test variables could potentially bias avoidance learning. Using an XGBoost decision tree model, we found a predictive relationship between exploratory/locomotor signatures during the Pre-Test and avoidance learning in the sCPA but not in the control object-conditioned cohort. Specifically, higher levels of exploration/locomotion were linked with reduced avoidance of the CS+ chamber. Finally, we tested the involvement of VTA GABA cells in avoidance learning in our protocol. Our data demonstrate that decreasing VTA inhibitory transmission during conditioning sessions is sufficient to abolish avoidance learning and its association with pre-existing exploratory/locomotor variables derived by our model. Our results establish a social defeat stress-inspired protocol for parsing contextual fear processes in a longitudinal and temporally discreet manner, confirming that VTA GABA cells are essential for the acquisition of this task.

Identifying vulnerable individuals based on pre-existing indices has been proposed as a powerful approach to potentially prevent and treat psychopathologies in a more personalized manner and it is currently an active research topic.^14,59–61^ However most studies in humans are able to do these assessments only *a posteriori*. Thus, research with animal models is indispensable to allow dissection of anxiety or other translatable traits as potential vulnerability factors prior to the trauma. Several behavioral tests using ethological conflicts have been developed and validated to study anxiety-related behaviors in rodents (for an overview please see Bailey and Crawley, 2009)^62^ and classifications based on composite anxiety-like behavior indices (*i.e.* comprising measurements from various anxiety tests) have been previously proposed.^14,63^ Previous studies have examined the role of individual differences such as reactivity to novelty and novelty seeking in fear learning and memory^64–68^ with higher locomotion and novelty seeking being associated with faster extinction in a fear conditioning protocol.^64^ However, if and how specific behavioral signatures influence specific stages of contextual social fear processing is not yet understood. Here, mice were classified based on their novelty-induced thigmotaxis, as this was the single variable that correlated with CS+ chamber avoidance in the sCPA. Next, using several readouts from the Pre-Test in an unbiased manner, we asked whether avoidance in the sCPA could be predicted by pre-existing behavioral traits. Among the Pre-Test features contributing to this broader predictive profile, rearing-related measures emerged as particularly prominent, with higher levels of rearing and locomotor activity being associated with reduced CS+ avoidance. Rearing has been previously shown to be context-and stress-sensitive and, moreover, it has been proposed as a potential measure of anxiety-like behavior in rodents.^69,70^ Specifically, Sturman et al., propose that distinguishing unsupported vs. supported rearing seems to tackle emotionality vs. activity-related components respectively.^69^ In our study, we did not differentiate between these two types of rearing. Nevertheless, as the sCPA apparatus chambers are quite compact, this involves potentially substantial levels of supported rearing to the walls, suggesting that the predictive contribution of rearing may, at least partly, reflect exploratory/locomotor tendencies rather than anxiety-like behavior per se. Despite the fact that the relationship between anxiety and exploration is quite complex, this association is documented in humans with high anxiety being associated with less exploration in foraging tasks.^71^ Here, we observed modest associations between thigmotaxis and the other predictive locomotor/exploratory features suggesting that anxiety-like behavior, novelty reactivity, locomotion and exploration are related but non-equivalent dimensions and that susceptibility to social avoidance can be potentially influenced by a multifactorial behavioral profile. One intriguing possibility is that, because exploration is critical for forming context-related associations,^23,24^ context-specific exploratory behavior might also facilitate memory updating. Future studies will need to explicitly address whether lower exploratory activity may lead to more persistent cue–context associations that are less amenable to extinction.

Freezing behavior in rodents is widely used in paradigms of fear conditioning to measure fear.^72^ However, other proxies are starting to emerge such as startle response,^73^ stress hormones, velocity,^27^ heart rate and flight responses.^74^ Our data highlight the need to adapt fear-related metrics as new paradigms of fear conditioning are developed. This requisite is showcased in the present study as when mice were allowed to explore the two adjacent chambers, freezing quantification revealed that they displayed more freezing in the CS-than CS+ chamber.

Our findings are in alignment with those from Tan et al. (2012) where increased freezing was observed in the unconditioned chamber after real-time place aversion conditioning induced by VTA GABA optogenetic activation.^37^ These findings could reflect a generalized anxiety state as previously proposed^37^ or a non-specific stress response associated with the general aversive experience. Our results also revealed higher locomotor activity when sCPA mice were in the CS+ chamber, as well as a pattern for faster exit and slower entry in the conditioned chamber. Taken together, our data highlights the importance of uncovering additional readouts and validating them as potential fear proxies when studying context processing.

Here, we show that VTA GABAergic transmission during conditioning sessions is essential for social fear learning in the sCPA. Our IHC experiments revealed a reduction of c-Fos^+^/TH^+^ cells in the sCPA vs. control condition, with c-Fos^+^/TH^-^ cells showing similar activity levels between groups. Previously, optogenetic activation of VTA GABA neurons was clearly shown to lead to real-time place aversion, an effect accompanied by inhibition of DA firing.^37^ Therefore, to indirectly manipulate the TH^+^ neurons, we selectively expressed an inhibitory Designer Receptors Exclusively Activated by Designer Drugs (DREADD) virus into the GABAergic population and assessed whether this would affect sCPA learning. Ideally, this question would be addressed by directly increasing the activity of VTA dopamine (DA) neurons during conditioning. However, because prolonged chemogenetic activation of midbrain DA neurons has been reported to induce dopaminergic neuron degeneration,^75^ we instead adopted an indirect strategy. Apart from GABAergic neurons, the TH^-^ population also includes glutamatergic neurons which are shown to play important roles in instrumental behaviors.^76^ Nevertheless, in the present study we found that selectively inhibiting the GABAergic population, abolished learning in the sCPA, suggesting that these neurons are necessary during conditioning. Along these lines, other studies have shown increased GCaMP6 fiber photometry signal in VTA Vgat neurons during social defeat when mice are being attacked. ^38^ In agreement with our data, inhibiting VTA GABA activity via chemogenetic manipulations abolished learning in an acute one-trial conditioned place aversion protocol.^31^

It is plausible that inhibiting VTA GABAergic neurons during conditioning leads to an attenuation of the aversive experience of social defeat. Our data measuring corticosterone after acute social defeat and within a CNO exposure window, show that hM4Di-injected individuals release less corticosterone than mCherry-injected controls, corroborating this interpretation. It has to be mentioned that inhibition of a specific GABAergic VTA population, namely Sst neurons, via chemogenetic approaches, led to opposite corticosterone concentration dynamics than what reported here.^38^ Cellular and projection-related heterogeneity of VTA GABA populations that could differentially be recruited by Vgat versus Sst-targeted approaches may contribute to the divergent physiological effects observed across studies.

An intriguing observation was that chemogenetic inhibition of VTA GABA neurons also abolished the predictive relationship between pre-conditioning behavioral state and subsequent avoidance learning. Importantly, the behavioral features used by the prediction model were acquired before conditioning and therefore before CNO administration and DREADD-mediated inhibition. Thus, the loss of predictive accuracy cannot be explained by changes in the baseline behavioral phenotype itself. Indeed, in control mCherry-injected animals, baseline behavioral features reliably predicted subsequent avoidance. One interesting possibility is that VTA GABA neurons could be playing a role bridging stable individual behavioral traits with inter-individual learning variability from aversive social experiences. Future experiments manipulating VTA GABA neurons after memory acquisition, or during context encoding, will be required to determine whether these neurons specifically gate the influence of pre-existing behavioral traits on learning or more generally support the acquisition of contextual aversive memories.

We cannot exclude that VTA GABA cells are exclusively important in social fear learning and not fear learning in general. Our protocol may serve as the basis for future characterization of other involved brain regions and circuits potentially by comparing them with readouts from electric shock protocols. Moreover, we do not suggest that this brain region and cell type are the sole critical contributors to learning in this task. It remains to be investigated whether the circuits underlying fear learning acquire social stress-related specialization in afferent or efferent pathways of the VTA.

One clear limitation of this study is that we included only male mice. Although we acknowledge that the field urgently needs to adapt this kind of protocols to females,^77^ according to our preliminary data aggression from CD-1 mice towards females was very low, even adapting according to previous protocols.^27,78,79^ However, given our finding that the amount of aggression received did not correlate with avoidance learning, it is plausible that low aggression levels could be sufficient to induce social place aversion in female mice. Elucidating this question remains our future goal as efforts to model social stress and violence with female rodents will offer significant translational benefits to researchers.

Although traditional fear conditioning protocols have contributed immensely in studying fear learning associations, the range of their ecological validity is debated.^25^ It is argued that developing and employing well-thought behavioral protocols is instrumental to advance our understanding of the brain.^80^ Thus, establishing replicable contextual learning protocols that exploit social defeat stress will allow us to delve further into multiple aspects of context-dependent fear processing, such as brain circuitry characterization, adaptation of protocols to female mice, and inter-individual differences. Here, we propose and characterize a longitudinal and robust contextual fear processing paradigm based on social stress and we offer evidence that exploratory/locomotor signatures and anxiety-related traits can shape specific acquisition and memory processes.

### Author contributions

LPP: Investigation, Visualization, Data Curation, Formal analysis, Validation, Methodology, Writing – review & editing.

AC: Conceptualization, Visualization, Data Curation, Validation, Methodology, Investigation, Formal analysis, Writing – review & editing.

PHPB: Methodology, Data Curation, Software, Formal analysis, Investigation.

CB: Investigation, Formal analysis.

OV: Investigation, Formal analysis. BA: Investigation, Formal analysis.

CP: Investigation, Formal analysis. ML: Investigation, Formal analysis.

ST: Conceptualization, Project administration, Funding acquisition, Resources, Visualization, Data Curation, Validation, Methodology, Supervision, Writing – original draft, Writing – review & editing.

## Acknowledgements

We thank Prof. Johannes Larsch, Dr. Clément Solié and Prof. Camilla Bellone for their thoughtful feedback on earlier drafts of the manuscript. We are especially grateful to Prof. Camilla Bellone for her continued support and advice throughout this work.

We would like to thank Prof. Anita Lüthi for kindly providing the first Vgat-cre breeding couples.

The laboratory of ST receives financial support from the Swiss National Science Foundation (SNSF), the Pierre Mercier Foundation, and the University of Lausanne (UNIL) Sophie Afenduli Foundation (accredited to L.P.P.).

AC is supported by the Swiss National Science Foundation and the David and Astrid Hagelén Foundation.

## Data availability statement

Code and processed datasets are available from the corresponding author upon reasonable request.

---

**Figure S1 -.**
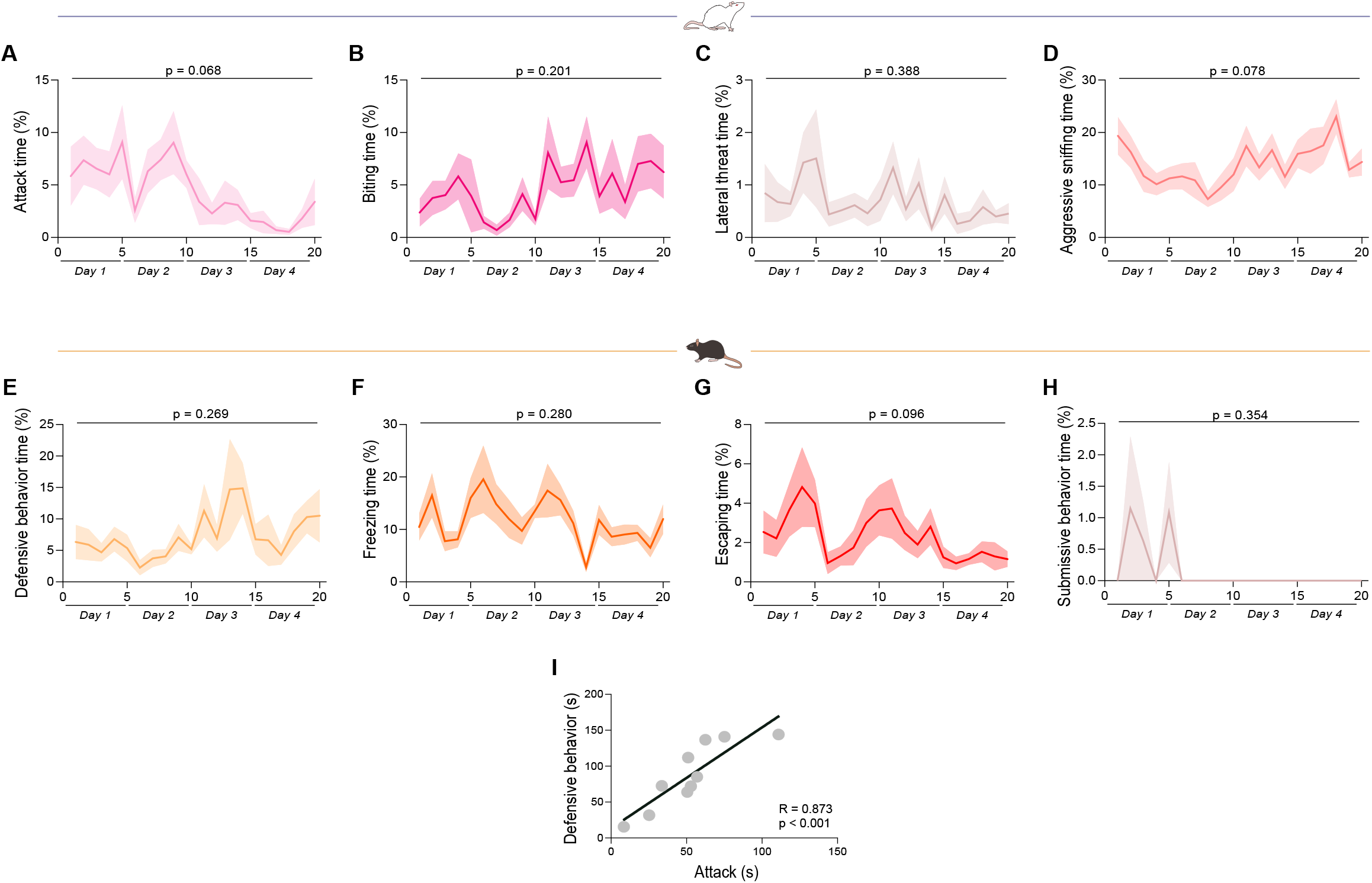
Behavioral characterization of conditioning sessions. **A-D**, Temporal dynamics of distinct CD-1 offensive behaviors during conditioning. Shaded areas represent mean ± SEM. A, Attack (RM one-way ANOVA with Greenhouse-Geisser correction): F(5.104, 45.933) = 2.209, p = 0.068; B, Biting (RM one-way ANOVA with Greenhouse-Geisser correction): F(4.746, 42.717) = 1.537, p = 0.201; C, Lateral threat (RM one-way ANOVA with Greenhouse-Geisser correction): F(3.366, 30.294) = 1.055, p = 0.388; D, Aggressive sniffing (RM one-way ANOVA with Greenhouse-Geisser correction): F(6.155, 55.399) = 2.010, p = 0.078. **E-H**, Temporal dynamics of distinct BL6 behaviors during conditioning. Shaded areas represent mean ± SEM. E, Defensive behavior (RM one-way ANOVA with Greenhouse-Geisser correction): F(3.951, 35.555) = 1.357, p = 0.269; F, Freezing (RM one-way ANOVA with Greenhouse-Geisser correction): F(5.279, 47.507) = 1.298, p = 0.280; G, Escaping (RM one-way ANOVA with Greenhouse-Geisser correction): F(4.172, 37.550) = 2.113, p = 0.096; H, Submissive behavior (RM one-way ANOVA with Greenhouse-Geisser correction): F(2.279, 20.515) = 1.115, p = 0.354. **I**, Relationship between attack received and defensive behavior exerted during conditioning. Each point represents one animal. Statistical test: Pearson’s correlation, R = 0.873, p < 0.001.

**Figure S2 -.**
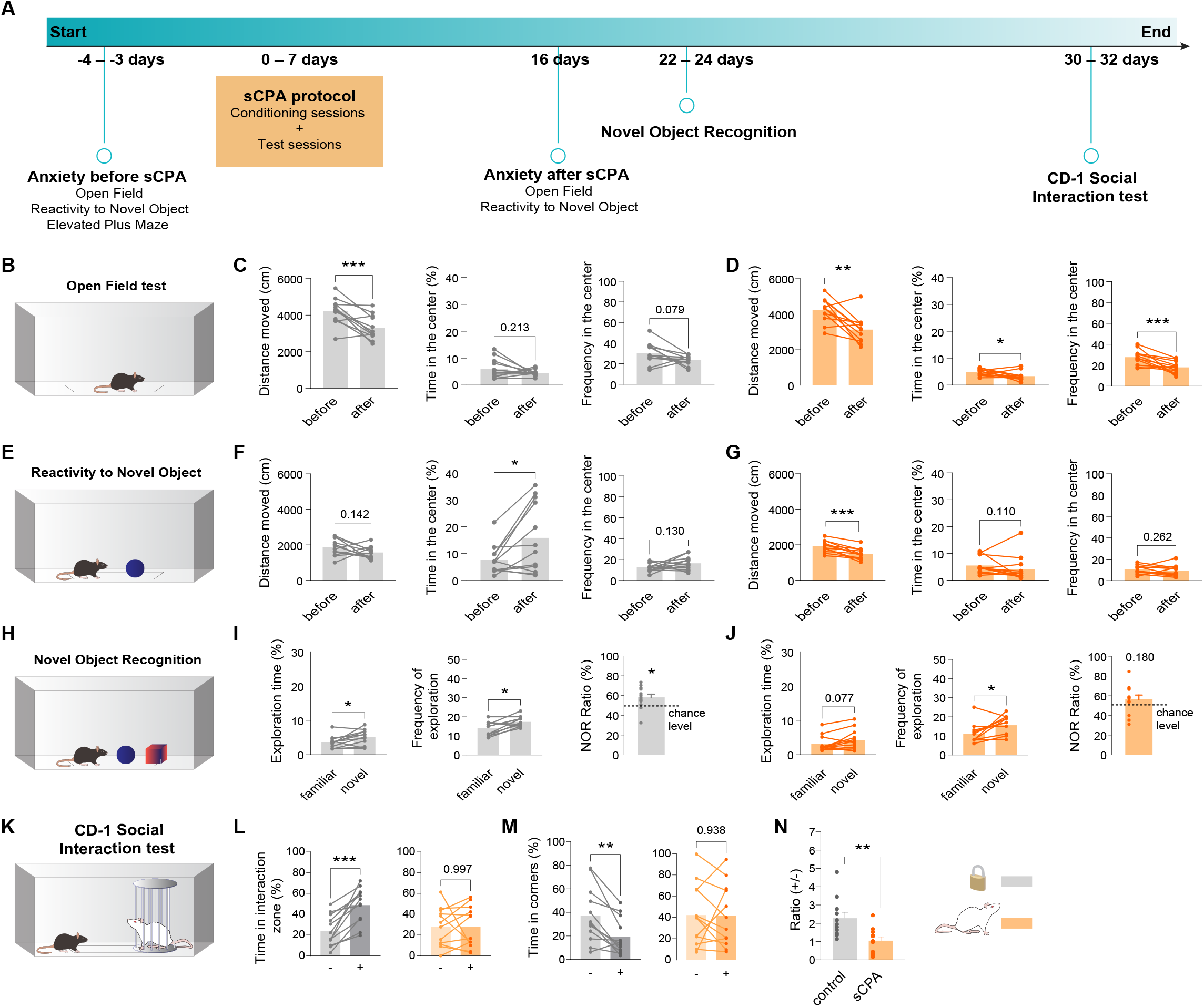
Effects of social conditioned place aversion on socioemotional and cognitive behavioral domains. **A**, Experimental timeline. A battery of anxiety-like tests (Open Field, reactivity to a Novel Object, Elevated Plus Maze tests) were performed before the social Conditioned Place Aversion paradigm to study the potential sources of inter-individual variability in stress responses. Another battery of anxiety-like tests (Open Field, reactivity to a Novel Object tests), as well as tests of novelty recognition and CD-1 Social Interaction were performed after the social conditioned place aversion paradigm to characterize stress effects on distinct behavioral domains. **B**, Schematic illustrating the Open Field test. **C**, Assessment of anxiety-like behavior with the Open Field task following object-based conditioning. Distance traveled (paired t-test): t(11) = 4.410, p = 0.001; Time in the center (paired t-test): t(11) = 1.321, p = 0.213; Frequency of center entries (Wilcoxon test): W = −45.000, p = 0.079. **D**, Assessment of anxiety-like behavior with the Open Field task following aversive social conditioning. Distance traveled (paired t-test): t(11) = 3.917, p = 0.002; Time in the center (Wilcoxon test): W = −52.000, p = 0.042; Frequency of center entries (Wilcoxon test): W = -75.000, p = 0.001. **E**, Schematic illustrating the reactivity to a Novel Object test. **F**, Assessment of exploratory behavior in an open field with the reactivity to a Novel Object test following object-based conditioning. Distance traveled (paired t-test): t(11) = 1.584, p = 0.142; Time in the center (Wilcoxon test): W = 52.000, p = 0.042; Frequency of center entries (Wilcoxon test): W = 35.000, p = 0.130. **G**, Assessment of exploratory behavior in an open field with the reactivity to a Novel Object test following aversive social conditioning. Distance traveled (paired t-test): t(11) = 5.970, p < 0.001; Time in the center (Wilcoxon test): W = −42.000, p = 0.110; Frequency of center entries (Wilcoxon test): W = −30.000, p = 0.262. **H**, Schematic illustrating the Novel Object Recognition test. **I**, Assessment of novelty recognition with the Novel Object Recognition test following object-based conditioning. Exploration time (paired t-test): t(11) = −2.624, p = 0.024; Frequency of exploration (Wilcoxon test): W = 54.000, p = 0.014; Novel Object Exploration Ratio (one-sample t-test): t(11) = 2.502, p = 0.029. **J**, Assessment of novelty recognition with the Novel Object Recognition test following aversive social conditioning. Exploration time (Wilcoxon test): W = 46.000, p = 0.077; Frequency of exploration (Wilcoxon test): W = 36.000, p = 0.031; Novel Object Exploration Ratio (one-sample t-test): t(11) = 1.433, p = 0.180. **K**, Schematic illustrating the CD-1 Social Interaction test. **L,** Time in the interaction zone in the absence (-) vs presence (+) of a novel CD-1 mouse following: Object-based conditioning (left) (paired t-test): t(11) = −5.815, p < 0.001; Aversive social conditioning (right) (paired t-test): t(11) = 0.004, p = 0.997. **M**, Time in the corners in the absence (-) vs presence (+) of a novel CD-1 mouse following: Object-based conditioning (left) (paired t-test): t(11) = 3.998, p = 0.002; Aversive social conditioning (right) (paired t-test): t(11) = 0.080, p = 0.938. **N**, CD-1 exploration Ratio (Time in the interaction zone (+) (s) / Time in the interaction zone (-) (s)) after object-based conditioning vs after aversive social conditioning (independent t-test): t(20) = −3.171, p = 0.005.

**Figure S3 -.**
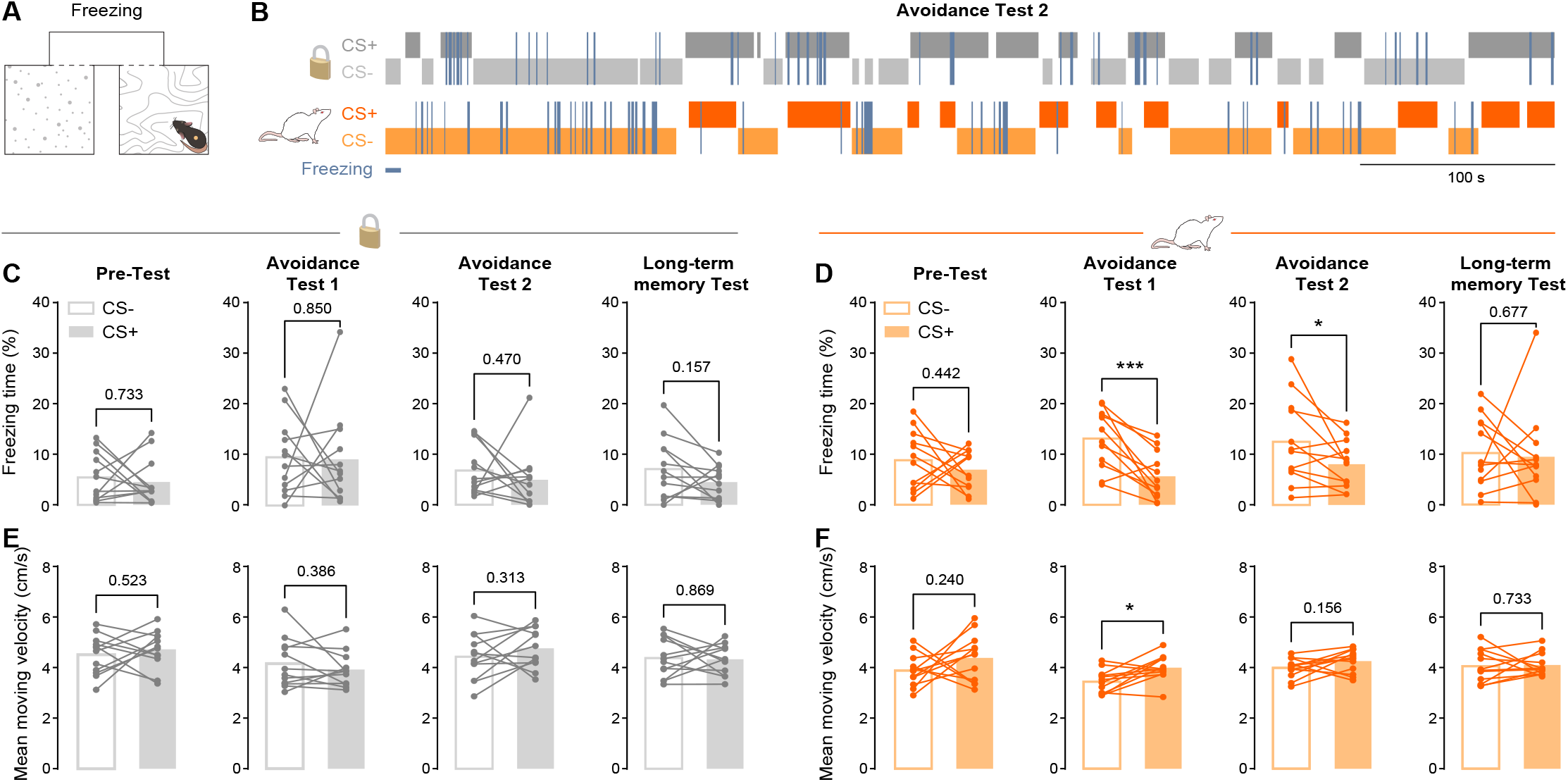
Freezing behavior and locomotor dynamics during expression of social conditioned place aversion. **A**, Schematic illustrating freezing behavior within the apparatus. **B**, Representative behavioral ethogram during Avoidance Test 2 showing chamber occupancy (CS+ and CS−) and freezing events for object-conditioned (top) and socially conditioned (bottom) animals. **C**, Freezing behavior during testing following object-based conditioning. Freezing time (%) was quantified separately in the CS− and CS+ chambers during the Pre-Test, Avoidance Test 1, Avoidance Test 2, and Long-term memory Test. Each line represents an individual animal. Pre-Test (Wilcoxon test): W = −10.000, p = 0.733; Avoidance Test 1 (Wilcoxon test): W = -6.000,p = 0.850; Avoidance Test 2 (Wilcoxon test): W = −20.000, p = 0.470; Long-term memory Test (paired t-test): t(11) = 1.518, p = 0.157. **D**, Freezing behavior during testing following aversive social conditioning. Freezing time (%) was quantified separately in the CS− and CS+ chambers during the Pre-Test, Avoidance Test 1, Avoidance Test 2, and Long-term memory Test. Each line represents an individual animal. Pre-Test (paired t-test): t(11) = 0.798, p = 0.442; Avoidance-Test 1 (paired t-test): t(11) = 4.440, p < 0.001; Avoidance-Test 2 (paired t-test): t(11) = 2.247, p = 0.046; Long-term memory Test (Wilcoxon test): W = −12.000, p = 0.677. **E**, Mean moving velocity per chamber during testing following object-based conditioning. Each line represents an individual animal. Pre-Test (paired t-test): t(11) = −0.659, p = 0.523; Avoidance Test 1 (paired t-test): t(11) = 0.903, p = 0.386; Avoidance Test 2 (paired t-test): t(11) = −1.057, p = 0.313; Long-term memory Test (paired t-test): t(11) = 0.168, p = 0.869. **F**, Mean moving velocity per chamber during testing following aversive social conditioning. Pre-Test (paired t-test): t(11) = −1.243, p = 0.240; Avoidance Test 1 (paired t-test): t(11) = −3.026, p = 0.012; Avoidance Test 2 (paired t-test): t(11) = −1.523, p = 0.156; Long-term memory Test (Wilcoxon test): W = 10.000, p = 0.733.

**Figure S4 -.**
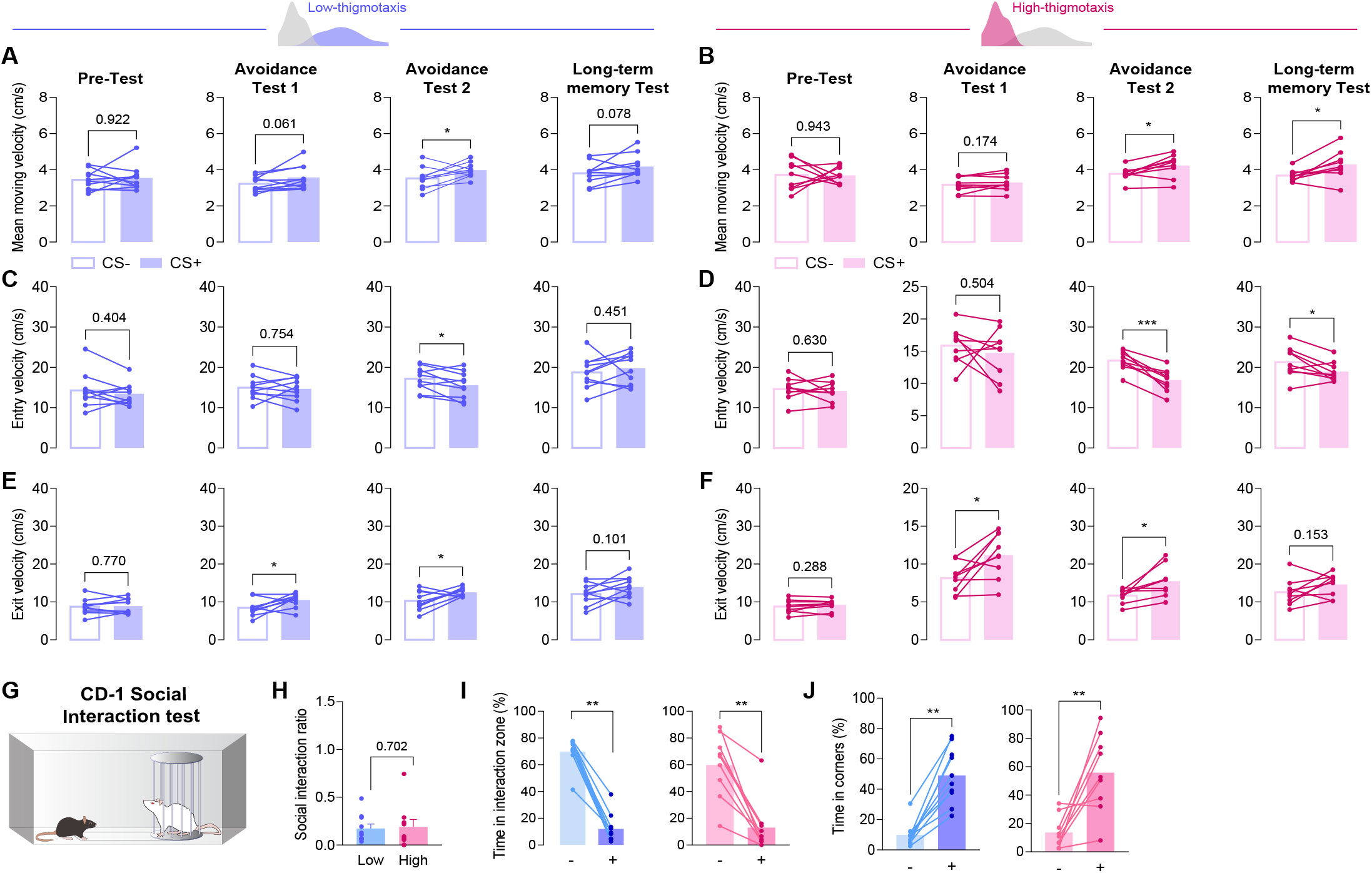
Locomotor dynamics during expression of social conditioned place aversion and social fear memory in novelty-induced high-and low-thigmotaxis animals. A, Mean moving velocity per chamber during testing following aversive social conditioning in low-thigmotaxis mice. Each line represents an individual animal. Pre-Test (Wilcoxon test): W = 3.000, p = 0.922; Avoidance Test 1 (paired t-test): t(9) = −2.145, p = 0.061; Avoidance Test 2 (paired t-test): t(9) = −2.324, p = 0.045; Long-term memory Test (paired t-test): t(9) = −1.992, p = 0.078. **B**, Mean moving velocity per chamber during testing following aversive social conditioning in high-thigmotaxis mice. Each line represents an individual animal. Pre-Test (paired t-test): t(8) = 0.073, p = 0.943; Avoidance Test 1 (paired t-test): t(8) = −1.492, p = 0.174; Avoidance Test 2 (paired t-test): t(8) = −3.210, p = 0.012; Long-term memory Test (paired t-test): t(8) = −2.884, p = 0.020. **C**, Mean peak velocity per chamber entry during testing following aversive social conditioning in low-thigmotaxis mice. Each line represents an individual animal. Pre-Test (paired t-test): t(9) = 0.875, p = 0.404; Avoidance Test 1 (paired t-test): t(9) = 0.323, p = 0.754; Avoidance-Test 2 (paired t-test): t(9) = 2.367, p = 0.042; Long-term memory Test (paired t-test): t(9) = −0.788, p = 0.451. **D**, Mean peak velocity per chamber entry during testing following aversive social conditioning in high-thigmotaxis mice. Each line represents an individual animal. Pre-Test (paired t-test): t(8) = 0.501, p = 0.630; Avoidance Test 1 (paired t-test): t(8) = 0.700, p = 0.504; Avoidance Test 2 (paired t-test): t(8) = 5.877, p < 0.001; Long-term memory Test (paired t-test): t(8) = 2.434, p = 0.041. **E**, Mean peak velocity per chamber exit during testing following aversive social conditioning in low-thigmotaxis mice. Each line represents an individual animal. Pre-Test (paired t-test): t(9) = −0.301, p = 0.770; Avoidance Test 1 (paired t-test): t(9) = −2.390, p = 0.041; Avoidance Test 2 (paired t-test): t(9) = −2.473, p = 0.035; Long-term memory Test (paired t-test): t(9) = −1.824, p = 0.101. **F**, Mean peak velocity per chamber exit during testing following aversive social conditioning in high-thigmotaxis mice. Each line represents an individual animal. Pre-Test (paired t-test): t(8) = −1.137, p = 0.288; Avoidance Test 1 (paired t-test): t(8) = −2.971, p = 0.018; Avoidance Test 2 (paired t-test): t(8) = −3.148, p = 0.014; Long-term memory Test (paired t-test): t(8) = −1.581, p = 0.153. **G**, Schematic illustrating the CD-1 Social Interaction test. **H**, CD-1 Social Interaction Ratio, defined as: Time in the interaction zone (+) (s) / Time in the interaction zone (-) (s), for low-vs high-thigmotaxis animals following aversive social conditioning (Mann Whitney test): U = 40.000, p = 0.702. **I**, Time in the interaction zone in the absence (-) vs presence (+) of a novel CD-1 mouse following aversive social conditioning for: Low-thigmotaxis mice (left) (Wilcoxon test): W = −55.000, p = 0.002; High-thigmotaxis mice (right) (Wilcoxon test): W = −45.000, p = 0.004. **J**, Time in the corners in the absence (-) vs presence (+) of a novel CD-1 mouse following aversive social conditioning for: Low-thigmotaxis mice (left) (Wilcoxon test): W = 55.000, p = 0.002; High-thigmotaxis mice (right) (paired t-test): t(8) = −3.922, p = 0.004.

**Figure S5 -.**
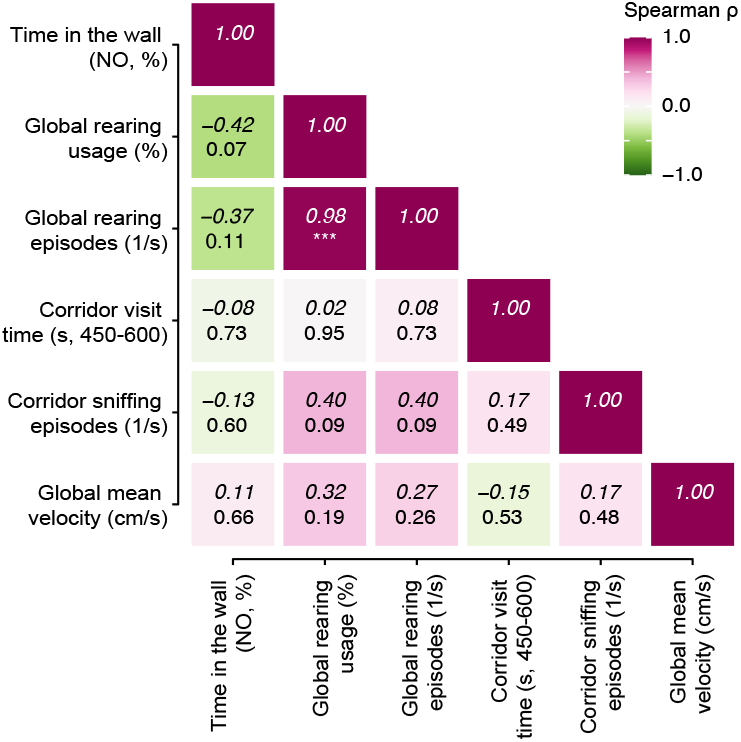
Relationship between novelty-induced thigmotaxis and Pre-Test predictive features. Spearman correlation matrix between novelty-induced thigmotaxis, measured as time spent in the wall zone during the Novel Object (NO) test and the main Pre-Test behavioral features contributing to the prediction of subsequent avoidance. Spearman’s correlation coefficients (ρ, top) and corresponding p-values (bottom) are reported for all pairwise comparisons.

**Figure S6 -.**
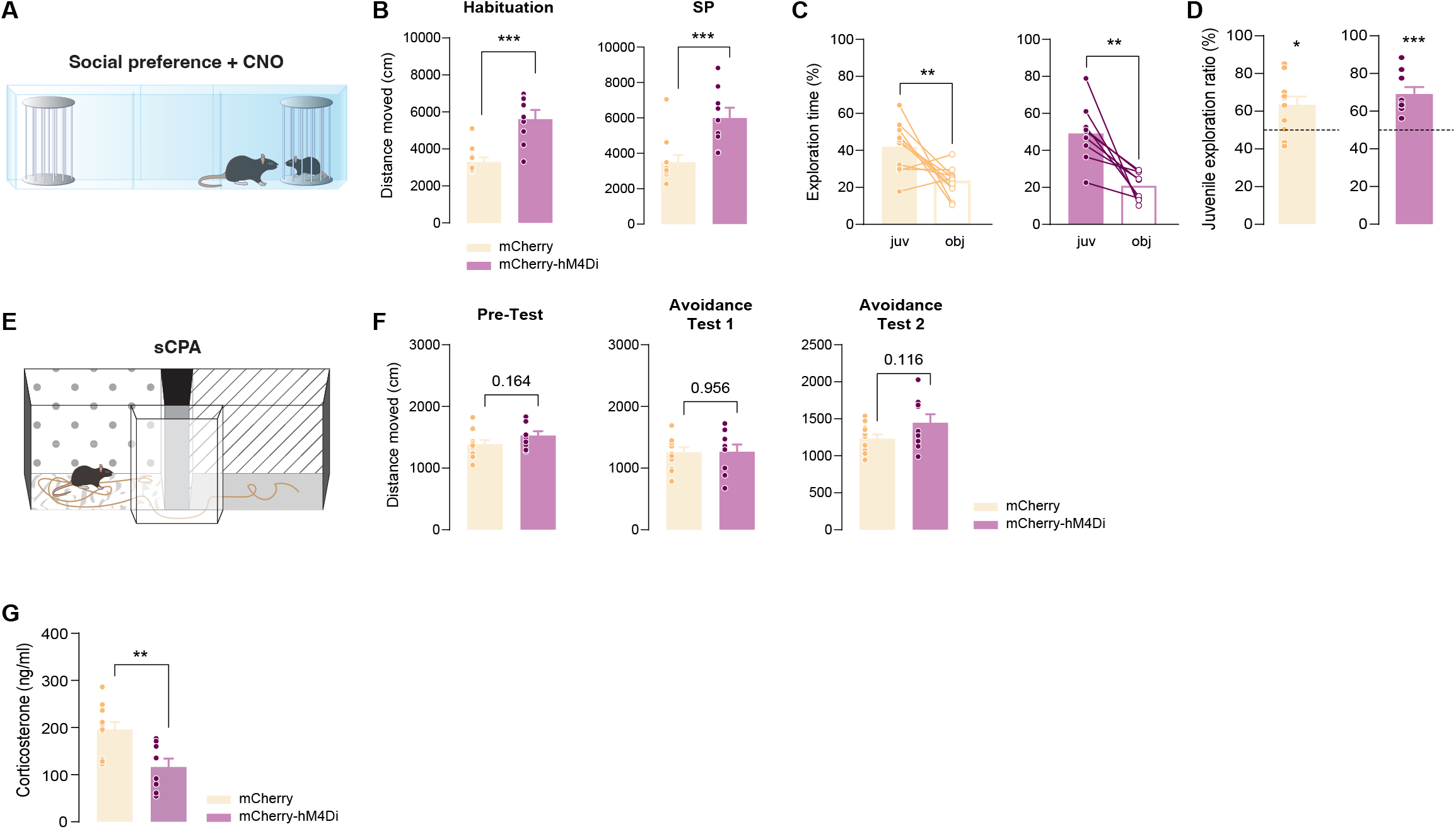
Inhibition of VTA^Vgat^ neurons during conditioning does not affect sociability, but rather impacts corticosterone reactivity. **A**, Schematic illustrating the Social Preference test. **B**, Distance moved in the absence (Habituation) and presence (SP) of a juvenile conspecific during the Social Preference test for the control mCherry and hM4Di-expressing mice. Habituation (Mann-Whitney test): U = 6.000, p <0.001; SP (Mann-Whitney test): U = 9.000, p = 0.001. **C**, Exploration time of a juvenile conspecific vs an object during the SP for the control mCherry and hM4Di-expressing mice: mCherry (left) (paired t-test): t(10) = 3.191, p = 0.010; mCherry-hM4Di (right) (paired t-test): t(8) = 4.287, p = 0.003. **D**, Juvenile exploration ratio, defined as: 100 ✕ Juvenile Exploration Time (s) / (Juvenile Exploration Time (s) + Object Exploration Time (s)), for the control mCherry and hM4Di-expressing mice: mCherry (left) (one-sample t-test against a 50% chance level): t(10) = 3.049, p = 0.012; mCherry-hM4Di (right) (one-sample t-test against a 50% chance level): t(8) = 5.275, p <0.001. **E**, Schematic illustrating the social Conditioned Place Aversion testing procedure. **F**, Distance moved during testing following for control mCherry and hM4Di-expressing mice: Pre-Test (independent t-test): t(19) = −1.447, p = 0.164; Avoidance Test 1 (independent t-test): t(19) = −0.056, p = 0.956; Avoidance Test 2 (Welch’s t-test): t(12.082) = −1.694, p = 0.116. **G**, Corticosterone reactivity to acute social defeat in control mCherry vs hM4Di-expressing mice: Statistical test: independent t-test: t(17) = 3.342, p = 0.004.

## Supplementary Tables legends

**Supplementary Table 1**: Contextual contribution of the training dataset.

**Supplementary Table 2:** Performance metrics.

**Supplementary Table 3:** Post-hoc filtering parameters.

**Supplementary Table 4:** Statistical analysis of pauses, sniffing, grooming, and rearing time (%) per chamber during the Pre-Test, Avoidance Test 1, Avoidance Test 2, and Long-term memory Test for the object-based conditioning (control) and aversive social conditioning (sCPA) groups.

**Supplementary Video 1:** Machine learning-labeled behavior examples after filtration and in different contexts.

