## Supplementary tables for "Individual behavioral signatures shape processing of context-dependent social fear"

| Training dataset video representation | | |
|---|---|---|
| **Arena** | **Number of videos** | **Number of videos with egocentric crop** |
| sCPA | 51 | 35 |
| Three chamber | 8 | 4 |
| EPM | 7 | 4 |
| Open-top cage and EPM | 1 | 0 |
| Open field | 22 | 15 |
| Skinner box | 5 | 0 |
| Total | 94 | 58 |

**Supplementary Table 1**: Contextual contribution of the training dataset.

| | **Precision** | **Recall** | **F1-score** | **Support** |
|---|---|---|---|---|
| Freezing | 0,906 | 0,977 | 0,94 | 1093 |
| Grooming | 0,971 | 9,929 | 0,95 | 1886 |
| Negative | 0,953 | 0,93 | 0,942 | 2946 |
| Rearing | 0,902 | 0,944 | 0,923 | 1268 |
| Sniffing | 0,89 | 0,893 | 0,891 | 1788 |
| Accuracy | | | 0,93 | 8981 |
| Macro avg | 0,924 | 0,935 | 0,929 | 8981 |
| Weighted avg | 0,931 | 0,93 | 93 | 8981 |

**Supplementary Table 2**: Performance metrics.

| Behavior | Confidence filter | Duration filter |
|---|---|---|
| Freezing | 50% | 0s |
| Grooming | 60% | 1,5s |
| Negative | 0 | 0s |
| Rearing | 70% | 0,8s |
| Sniffing | 30% | 0s |

**Supplementary Table 3**: Post-hoc filtering parameters.

| | | Object-based conditioning (control) | | | | | | | | | Aversive social conditioning (sCPA) | | | | | | | | |
|---|---|---|---|---|---|---|---|---|---|---|---|---|---|---|---|---|---|---|---|
| | | Mean (CS-) | SEM (CS-) | SD (CS-) | Mean (CS+) | SEM (CS+) | SD (CS+) | Test | Test statistic | p-value | Mean (CS-) | SEM (CS-) | SD (CS-) | Mean (CS+) | SEM (CS+) | SD (CS+) | Test | Test statistic | p-value |
| **Pauses (%)** | Pre-Test | 10.635 | 3.473 | 12.030 | 6.677 | 2.231 | 7.728 | Wilcoxon | -12.000 | 0.677 | 13.205 | 3.339 | 11.566 | 9.662 | 2.240 | 7.761 | Wilcoxon | -12.000 | 0.677 |
| | Avoidance Test 1 | 28.059 | 6.768 | 23.444 | 23.742 | 4.924 | 17.057 | paired t-test | 0.429 | 0.677 | 30.090 | 3.921 | 13.583 | 16.121 | 4.868 | 16.862 | Wilcoxon | -68.000 | 0.005* |
| | Avoidance Test 2 | 18.961 | 4.231 | 14.658 | 11.692 | 2.877 | 9.965 | paired t-test | 1.342 | 0.206 | 25.447 | 4.077 | 14.122 | 15.411 | 3.473 | 12.031 | paired t-test | 1.868 | 0.089 |
| | Long-term memory Test | 23.094 | 4.666 | 16.163 | 14.625 | 4.321 | 14.968 | Wilcoxon | -34.000 | 0.204 | 23.013 | 6.074 | 21.041 | 21.386 | 5.865 | 20.316 | paired t-test | 0.165 | 0.872 |
| **Sniffing (%)** | Pre-Test | 33.180 | 2.259 | 7.825 | 32.962 | 3.023 | 10.474 | paired t-test | 0.063 | 0.951 | 36.294 | 2.446 | 8.472 | 31.918 | 2.892 | 10.018 | paired t-test | 1.303 | 0.219 |
| | Avoidance Test 1 | 30.724 | 2.305 | 7.986 | 31.486 | 2.822 | 9.774 | paired t-test | -0.226 | 0.825 | 28.840 | 2.729 | 9.452 | 28.744 | 2.290 | 7.934 | paired t-test | 0.024 | 0.981 |
| | Avoidance Test 2 | 27.538 | 3.048 | 10.560 | 26.384 | 2.176 | 7.539 | paired t-test | 0.270 | 0.792 | 28.604 | 2.025 | 7.014 | 26.966 | 2.405 | 8.330 | Wilcoxon | -10.000 | 0.733 |
| | Long-term memory Test | 27.071 | 1.974 | 6.838 | 24.990 | 1.714 | 5.938 | paired t-test | 0.671 | 0.516 | 28.092 | 1.950 | 6.753 | 22.828 | 2.144 | 7.426 | Wilcoxon | -48.000 | 0.064 |
| **Grooming (%)** | Pre-Test | 3.800 | 1.443 | 4.998 | 2.839 | 1.118 | 3.874 | Wilcoxon | -11.000 | 0.625 | 3.323 | 1.697 | 5.878 | 2.011 | 0.661 | 2.290 | Wilcoxon | 4.000 | 0.898 |
| | Avoidance Test 1 | 3.932 | 1.526 | 5.285 | 3.278 | 1.000 | 3.465 | Wilcoxon | -2.000 | 0.970 | 5.537 | 1.860 | 6.443 | 4.877 | 3.137 | 10.868 | Wilcoxon | -22.000 | 0.424 |
| | Avoidance Test 2 | 5.886 | 2.433 | 8.427 | 2.440 | 0.816 | 2.828 | Wilcoxon | -18.000 | 0.519 | 6.228 | 2.067 | 7.160 | 3.394 | 1.541 | 5.337 | Wilcoxon | -22.000 | 0.424 |
| | Long-term memory Test | 5.493 | 1.387 | 4.804 | 4.001 | 1.328 | 4.600 | Wilcoxon | -22.000 | 0.424 | 4.773 | 1.927 | 6.676 | 4.024 | 1.741 | 6.032 | Wilcoxon | -2.000 | 0.966 |
| **Rearing (%)** | Pre-Test | 8.257 | 1.306 | 4.524 | 7.795 | 1.411 | 4.888 | paired t-test | 0.230 | 0.823 | 7.011 | 1.388 | 4.807 | 6.412 | 0.637 | 2.206 | Wilcoxon | 8.000 | 0.791 |
| | Avoidance Test 1 | 8.538 | 1.928 | 6.678 | 6.167 | 1.129 | 3.910 | Wilcoxon | -24.000 | 0.380 | 7.020 | 1.100 | 3.809 | 9.567 | 1.409 | 4.881 | paired t-test | -1.266 | 0.232 |
| | Avoidance Test 2 | 10.256 | 2.148 | 7.442 | 11.080 | 1.378 | 4.775 | Wilcoxon | 16.000 | 0.569 | 7.123 | 1.139 | 3.947 | 8.984 | 1.533 | 5.311 | paired t-test | -1.128 | 0.283 |
| | Long-term memory Test | 8.018 | 1.570 | 5.440 | 9.104 | 1.300 | 4.503 | paired t-test | -0.462 | 0.653 | 9.672 | 2.212 | 7.662 | 9.457 | 1.856 | 6.428 | Wilcoxon | 26.000 | 0.339 |

**Supplementary Table 4**: Statistical analysis of pauses, sniffing, grooming, and rearing time (%) per chamber during the Pre-Test, Avoidance Test 1, Avoidance Test 2, and Long-term memory Test for the object-based conditioning (control) and aversive social conditioning (sCPA) groups.
